# *Endozoicomonas*-chlamydiae interactions in cell-associated microbial aggregates of the coral *Pocillopora acuta*

**DOI:** 10.1101/2022.11.28.517745

**Authors:** Justin Maire, Kshitij Tandon, Astrid Collingro, Allison van de Meene, Katarina Damjanovic, Cecilie Ravn Gøtze, Sophie Stephenson, Gayle K. Philip, Matthias Horn, Neal E. Cantin, Linda L. Blackall, Madeleine J. H. van Oppen

**Affiliations:** School of BioSciences, The University of Melbourne, Parkville, VIC, 3010, Australia; Centre for Microbiology and Environmental Systems Science, University of Vienna, Vienna, 1030, Austria; Australian Institute of Marine Science, PMB No 3, Townsville, 4810, QLD, Australia; Melbourne Bioinformatics, The University of Melbourne, Parkville, VIC, 3010, Australia

**Keywords:** microbiome, *Endozoicomonas*, CAMA, endosymbiont, coral, chlamydiae

## Abstract

Corals are associated with a variety of bacteria, which occur in the surface mucus layer, gastrovascular cavity, skeleton, and tissues. Some tissue-associated bacteria form clusters, termed cell-associated microbial aggregates (CAMAs), which are poorly studied. Here, we provide a comprehensive characterization of CAMAs in the coral *Pocillopora acuta*. Combining imaging techniques, laser capture microdissection, and amplicon and metagenome sequencing we show that CAMAs: (i) are located in the tentacle tips and may be intracellular; (ii) contain *Endozoicomonas, Kistimonas* (both Gammaproteobacteria), and *Simkania* (Chlamydiota) bacteria; (iii) *Endozoicomonas* may provide vitamins to its host and use secretion systems and/or pili for colonization and aggregation; (iv) *Endozoicomonas* and *Simkania* occur in distinct, but adjacent, CAMAs; (v) *Simkania* may rely on acetate and heme provided by neighboring *Endozoicomonas*. Our study provides detailed insight into coral endosymbionts, which will guide the assessment of their suitability for probiotic approaches to mitigate coral bleaching.

## Introduction

Coral reefs are among the most important ecosystems on the planet because they constitute hotspots for biodiversity (*1*), and provide many goods and services, including coastal protection and important industries such as tourism and fisheries (*2*). Scleractinian corals are foundation members of coral reefs, as their skeleton forms the reefs’ three-dimensional structure that provides habitats for many reef-dwelling organisms and because they are the main primary producers on the reef. Corals rely on a wide range of microorganisms for their survival, including protists, bacteria, archaea, fungi and viruses (*3–5*). Photosynthetic dinoflagellates of the Symbiodiniaceae family are by far the most studied coral-associated microorganisms. They are vital symbionts because they provide their hosts with most of their energy through the translocation of photosynthates (*6*). Some coral-associated bacteria also have roles that are critical to the coral host, such as protection against pathogens (*7*), and cycling of nitrogen and sulfur (*8–10*).

Bacteria can colonize all microhabitats in corals and are most abundant and diverse in the mucus and skeleton (*3, 4*). Tissue-associated bacteria are less diverse (*9, 11*) and some are known to form large, dense clusters termed cell-associated microbial aggregates (CAMAs). Initially described in the early 1980s, they were thought to be linked to coral disease in *Acropora palmata* (*12, 13*). CAMAs have since been described in numerous cnidarian species, including hard corals, soft corals, and anemones, regardless of disease status, and are particularly common in the genera *Acropora, Pocillopora, Porites, Platygyra* and *Stylophora* (*14–23*). Apart from location and distribution within the coral host, knowledge about CAMAs is scarce, with a handful of studies applying taxa-specific histological techniques to identify bacteria residing in CAMAs (*16, 19, 23, 24*), and only one recent study applying culture-independent genomics to assess their functional potential (*21*). These studies have shown that bacteria residing in CAMAs often belong to the *Endozoicomonas* genus, a widespread coral symbiont (*24*). Several genomes of *Endozoicomonas* cultured from corals have been sequenced (*10, 25, 26*), but the location of these strains within the coral host and their ability to form CAMAs was not assessed. Hence, a holistic understanding of coral-associated CAMAs is still lacking, despite the huge progress made in coral microbiome research in the past couple of decades.

Here, we provide a detailed characterization of CAMAs in the coral *Pocillopora acuta*, including their location, subcellular structure, transmission, community composition, and functional potential (Figure S1, Table S1). *Pocillopora acuta* is an asexual brooder and sexual broadcast spawner coral (*27*). It acquires bacteria both vertically, from the mother colony, and horizontally, from the environment (*28*). Associated bacterial communities are often dominated by *Endozoicomonas* (*28, 29*), and CAMAs have been detected in asexually produced larvae, suggesting their vertical transmission in this species (*28*). Using a combination of fluorescence *in situ* hybridization (FISH), confocal laser scanning microscopy (CLSM), and transmission electron microscopy (TEM), we show that CAMAs are located in the epidermal layer in tentacle tips of *P. acuta* polyps and are likely intracellular. CAMAs were precisely excised with laser capture microdissection (LCM) and community composition analysis showed they contain bacteria of the families Endozoicomonadaceae and Simkaniaceae, a member of the Chlamydiota phylum. Metagenomic analyses suggest Endozoicomonadaceae may be beneficial to its coral host, but also to Simkaniaceae present in adjacent CAMAs.

## Results and discussion

### CAMAs are located in *Pocillopora acuta’*s tentacles

*Pocillopora acuta* colonies were sampled from the field, reared in captivity over a three-year period, and asexually produced two successive generations of corals (Figure S2A). The two latter generations were used for this study and are hereafter referred to as generations F1 and F2. To investigate the location of CAMAs within *P. acuta* polyps, we first conducted whole-mount FISH with a universal bacterial probe (EUB338-mix). CLSM images showed that CAMAs are present at the tip of the tentacles (Figure 1A-C), but not anywhere else in the polyp. This was the case for all three genotypes we sampled (Figure S2A), C2_12, R2_8, and F1_6. Nematocysts, present in the tentacles, displayed significant non-specific binding, as evidenced by the simultaneous use of an antisense probe (Figure 1A). FISH and CLSM performed on polyp sections yielded the same pattern (Figure 1D-E). Analysis of adult offspring (F2 generation) samples also revealed a multitude of CAMAs at the tip of the tentacles (Figure 1F-G). *In situ* location in other cnidarians has failed to show any consistency across species: CAMAs have been found in the tentacles of *Exaiptasia diaphana* (*18, 20*) and *Stylophora pistillata* (*21*), the mesenteries of *Acropora formosa* and *A. palmata* (*12, 14*), and throughout all tissue types in *Acropora hyacinthus* (*22*). We observed between 0 and 10 CAMAs per polyp in *P. acuta*, and they measured between 20 and 80 μm in diameter, which is similar to other coral species (*21, 23*). CAMAs were exclusively seen in the epidermis and are thus not in contact with Symbiodiniaceae cells (Figure 1E), which exist in the gastroderm. Therefore, CAMA bacteria are more likely to interact with the coral animal than with the Symbiodiniaceae in *P. acuta*. CAMAs have only been detected in the epidermis in *E. diaphana* (*18, 20*) and *S. pistillata* (*19*), while they colonize the gastrodermis in *Pocillopora verrucosa* (*19*), *S. pistillata* (*14, 15, 19*), *Acropora aspera* (*15*) and *A. formosa* (*14*). This variability in location suggests that CAMAs may play distinct roles across coral holobionts. For example, gastrodermal CAMAs may interact with Symbiodiniaceae, while CAMAs in tentacles may bear defensive functions. Additionally, distinct infection mechanisms may exist and allow different bacterial taxa to colonize and aggregate in different tissues, *e.g*. tentacles may be a privileged site of contact between the coral and environmental bacteria.

**Figure 1:**
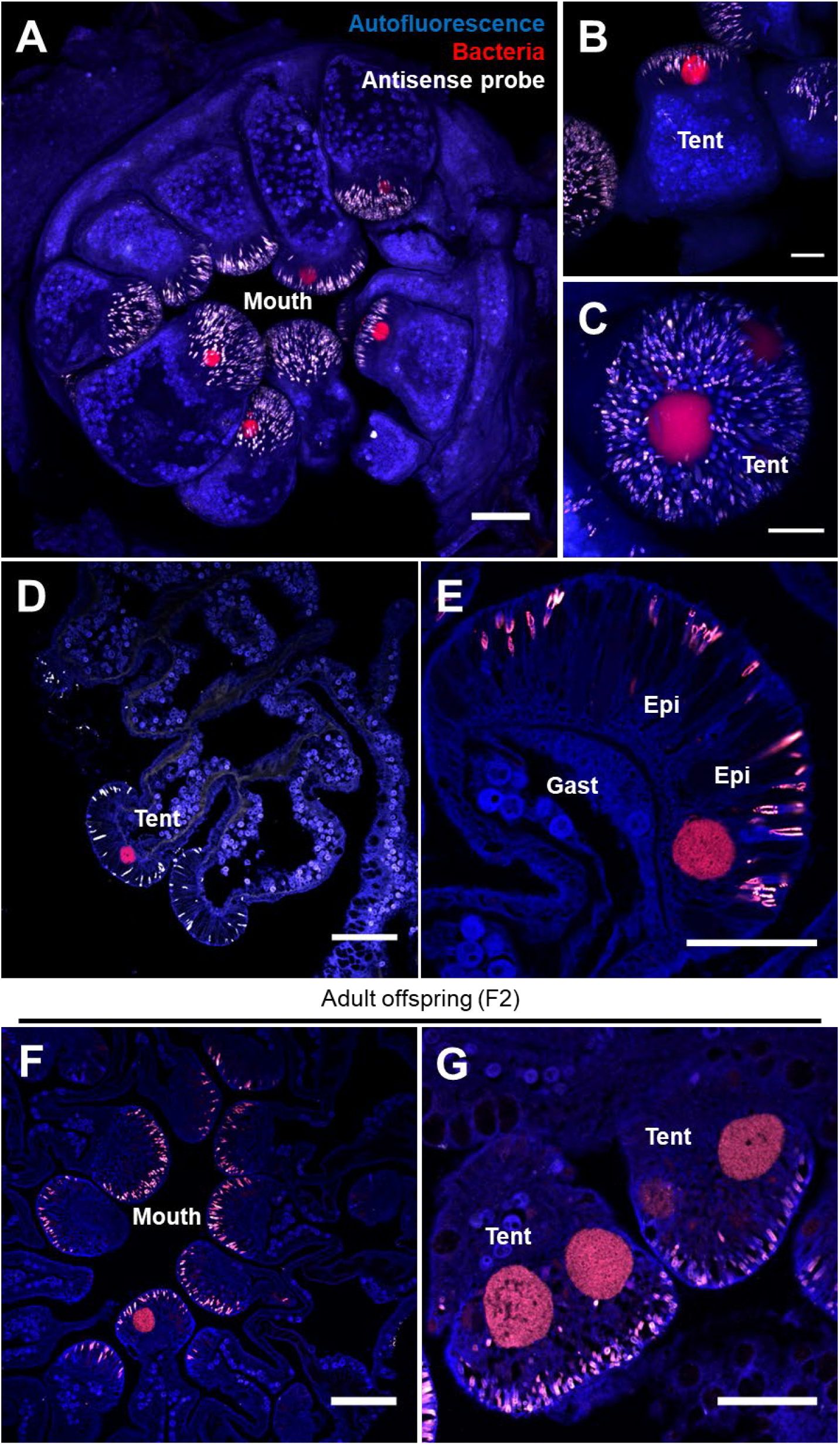
CAMAs are located in the epidermis of *P. acuta’*s tentacles. A-C: CAMA location by whole-mount FISH and CLSM in a whole adult polyp (A, maximum projection of a z-stack) and tentacles (B, C). D-G: CAMA location by FISH on sectioned adult polyps. Genotypes: R2_8 (A, B); F1_6 (C, E, F); C2_12 (D, G). Generation: F1 (A-E), F2 (F, G). Blue: autofluorescence; red: EUB338-mix probe (all bacteria); white: non-EUB probe (negative control). Gast: Gastrodermis; Epi: Epidermis; Tent: Tentacle. Scale bars: 100 μm for A, D, F; 50 μm for B, C, E, G.

While CAMAs are consistently found within cnidarian tissues, whether they are intra- or extracellular varies; they are intracellular in *E. diaphana* (*20*), extracellular in *A. palmata* (*13*), and both intra- and extracellular in *Porites compressa* (*23*). TEM images revealed that CAMAs are densely packed with bacteria (Figure 2A-B), which all show similar morphology and a highly visible nucleoid region (Figure 2C-D). TEM images in a previous study also showed the presence of nucleoid regions in CAMA bacteria (*20*). We observed a membrane encasing the CAMAs (Figures 2C-D and S3). This membrane resembled other coral cell membranes (Figure 2C, S3A and S3F; see for example the membrane of a cnidocyte, surrounding a nematocyst in Figure S3E) and was composed of lipid bilayers (Figure S3D. This suggests that CAMAs are intracellular in *P. acuta*. Because no eukaryotic organelle was observed inside CAMAs, it is possible that CAMAs are encased inside a vacuole, inside a coral cell. This is supported by the presence of eukaryotic content near CAMAs (*e.g.*, Golgi apparatus in Figure 2D), and the presence of a second membrane seemingly surrounding the CAMA membrane (Figures 2C, S3A, S3B, S3C, and S3F), and sometimes fusing with the CAMA membrane (Figures S3A and S3C). Alternatively, CAMAs may be extracellular, *i.e*. in between coral cells, and contained in a membrane of prokaryotic origin. DAPI staining combined with FISH showed the presence of one or more nuclei within CAMAs (Figure 2E-F), which is consistent with an intracellular nature. We also observed CAMAs that seemed to be disintegrating, with bacteria of a similar morphology seen outside the CAMAs (Figure 2F). This suggests that CAMAs are dynamic within the coral holobiont and may be able to form and deform after colonization of the animal. The mechanisms by which these bacteria aggregate, and whether the process is controlled by the host, bacteria, or both, require further investigation.

**Figure 2:**
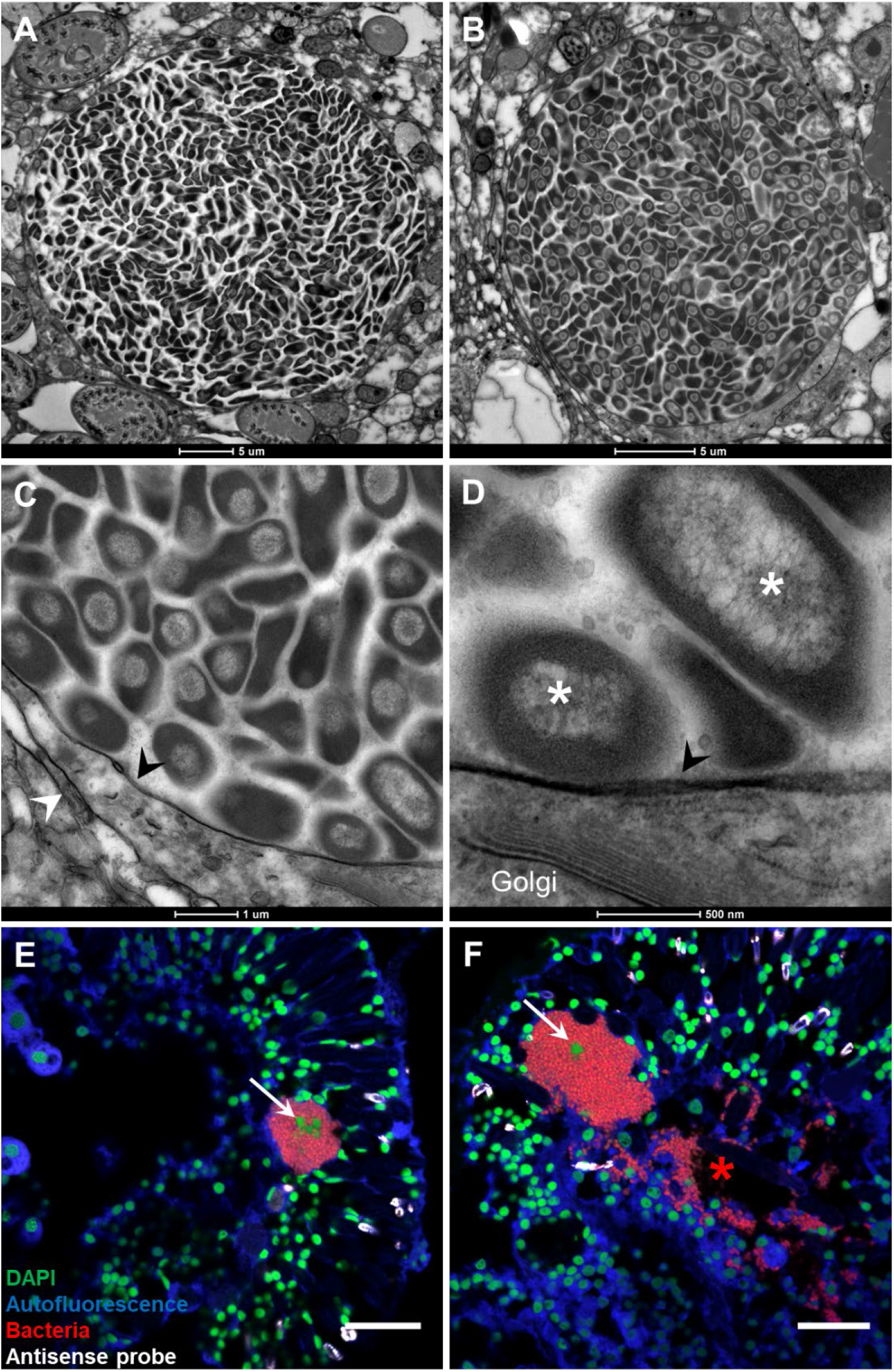
CAMAs may be intracellular. A-D: Subcellular structure of CAMAs observed in TEM. Black arrowheads point at a possible membrane surrounding the CAMAs. White arrowheads point at possible coral cell membranes. White asterisks show dense nucleoid regions of bacteria. E, F: Combined FISH and DAPI staining observed by CLSM on sectioned adult polyps. White arrows point at host nuclei inside CAMAs. The red asterisk highlights a zone where bacteria are outside a CAMA. All images are from F1 generation samples. Genotype: C2_12 (A-E); F1_6 (F). Blue: autofluorescence; red: EUB338-mix probe (all bacteria); white: non-EUB probe (negative control), green: DAPI. Scale bars: 20 μm for E, F.

### Endozoicomonadaceae and Simkaniaceae are the main CAMA members

Using laser capture microdissection (LCM), we specifically sampled CAMAs from two successive generations (F1 and F2, Figure S2A) of adult coral colonies belonging to the single F1_6 genotype (Figure S4A), that were raised entirely under controlled experimental conditions. 16S rRNA gene metabarcoding was performed to investigate the taxonomy of the bacteria contained in CAMAs. Sequencing statistics for this and the following metabarcoding experiments are summarized in Table S2. Fourteen ASVs were detected, although only five had a relative abundance above 0.1% in at least one sample (Table S3A). Four of those were assigned to the *Endozoicomonas* genus, were present in all samples, and amounted to more than 95% of all reads (Figure 3A). ASVs 01, 02, 03 showed >99% identity with each other (Figure S5), suggesting that they might belong to one single *Endozoicomonas* strain, while ASV04 was more distant (~96% identity to ASVs 01, 02, and 03) and is likely a separate strain. *Endozoicomonas* is widespread in marine invertebrates, including corals (*19, 24*), and sometimes dominates assemblages of coral-associated bacterial communities (*16, 21, 28, 30*).

**Figure 3:**
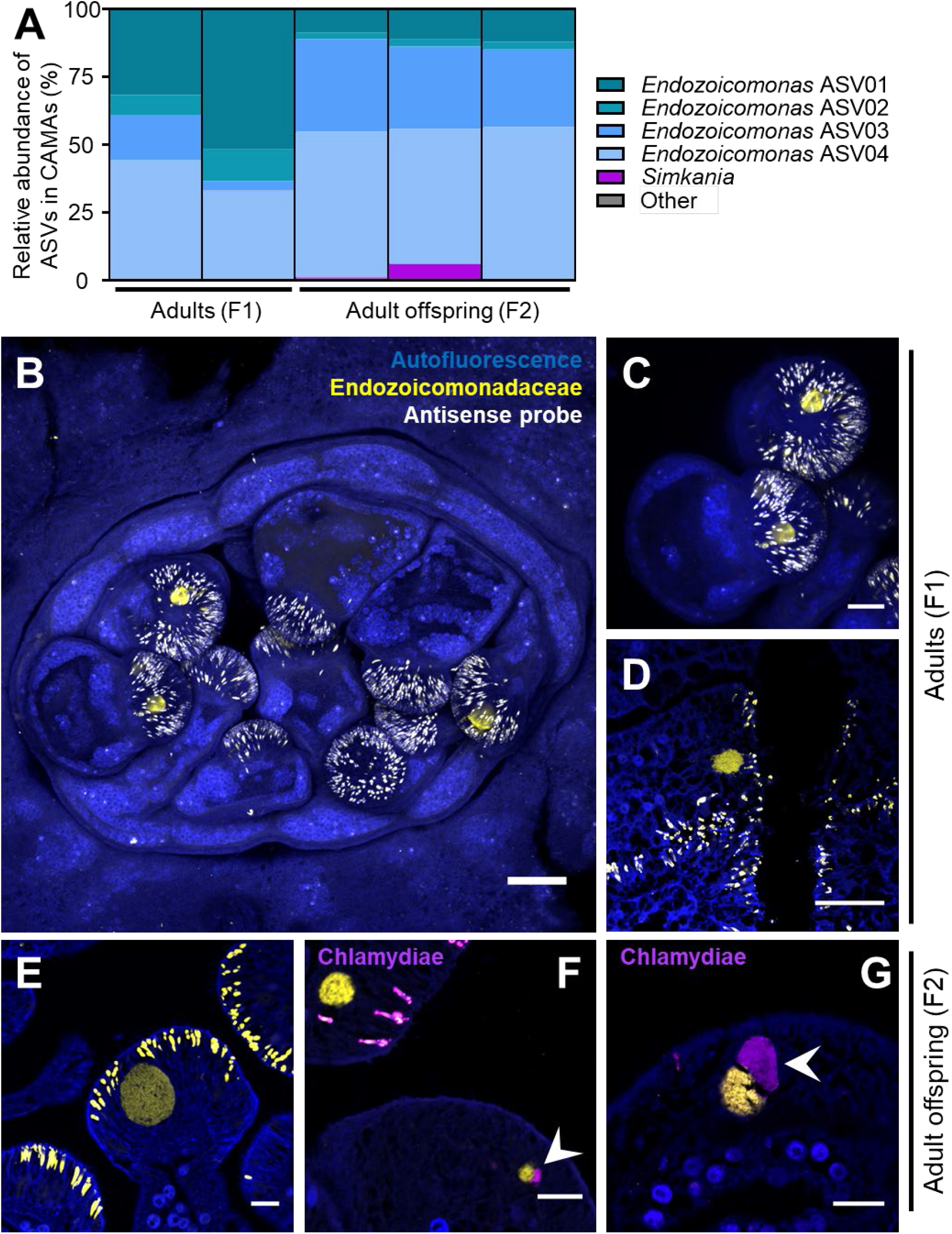
*Endozoicomonas* and *Simkania* are the two major taxa in CAMAs of the F1_6 genotype. A: Relative abundance of bacterial ASVs in CAMAs of adult polyps (F1_6 genotype, F1 and F2 generation) isolated by LCM. Each bar is a single sample. ASVs classified as ‘others’ represent less than 0.1% of the communities and cannot be seen in the figure. Detailed composition is available in Table S3. B-C: Location of *Endozoicomonas* in CAMAs by whole-mount FISH in a whole adult polyp (B, maximum projection of a z-stack) and tentacle (C). D-G: Location of *Endozoicomonas* and *Simkania* in CAMAs by FISH on sectioned adult polyps. White arrowheads point at *Simkania* CAMAs (purple), adjacent to *Endozoicomonas* CAMAs (yellow). All photos are from the F1_6 genotype. Generation: F1 (B-D), F2 (E-G). Blue: autofluorescence; yellow: End663 probe (Endozoicomonadaceae); magenta: Chls523 probe (chlamydiae). Scale bars: 100 μm for B; 50 μm for C, D; 25 μm for E, F.

FISH using a previously published *Endozoicomonas* probe (*16*) confirmed that *Endozoicomonas* are present in the CAMAs (Figure 3B-E). Simultaneous use of the *Endozoicomonas* probe with a universal bacterial probe showed complete co-localization (Figure S6A), confirming that *Endozoicomonas* are the main, and likely only, bacteria making up these CAMAs. FISH using the same probe has previously shown the presence of *Endozoicomonas* in CAMAs of *S. pistillata* and *P. verrucosa* (*16, 19, 24*), suggesting that *Endozoicomonas* commonly forms CAMAs in a range of coral species. Endozoicomonadaceae also form aggregates in the tissues of ascidians and fish (*31, 32*), and inside the nuclei of mussel cells (*33*), suggesting that a conserved mechanism across this bacterial family may allow them to form aggregates inside animal hosts. Recently, Wada et al. (2022) also used LCM to sample single CAMAs and confirmed with 16S rRNA gene metabarcoding the presence of several *Endozoicomonas* strains within individual CAMAs of *S. pistillata* (*21*). Because we pooled many CAMAs in our samples and used a genus-wide *Endozoicomonas* FISH probe, we cannot say whether several strains co-occur in a single CAMA in *P. acuta*. The co-occurrence of two or more *Endozoicomonas* strains in CAMAs is puzzling and suggests they might show a degree of metabolic complementation or dependence on one another. It also raises the question of strain compatibility and whether all *Endozoicomonas* strains can co-exist within a CAMA.

The last ASV with a relative abundance over 0.5% was assigned to the *Simkania* genus (Figure 3A) and was present in just two samples, both belonging to the F2 generation. The Simkaniaceae family belongs to the Chlamydiota phylum, which are obligate intracellular bacteria infecting a vast array of animals and thriving as symbionts of protists in diverse environments (*34*). Simkaniaceae ASVs are often detected at high relative abundances in coral samples (*21, 30, 35*), as well as in Symbiodiniaceae cultures (*36, 37*), although their roles, as well as the cells they infect, remain uninvestigated. In the present study, FISH using a previously published chlamydiae probe (*38*), simultaneously with the *Endozoicomonas* probe, showed *Simkania* inclusions are distinct from, but always adjacent to *Endozoicomonas* CAMAs in *P. acuta* (Figure 3F-G). Use of the chlamydiae probe combined with the universal bacterial probe confirmed *Simkania* occur alone within their inclusions (Figure S6B). While this is the first evidence of *Simkania* inclusions in corals, use of the Gimenez stain has previously suggested the presence of *Chlamydia-* or *Rickettsia-like* bacteria inclusions in *Acropora muricata* (*23*). Because chlamydiae are intracellular bacteria, the presence of *Simkania* inclusions reinforces the hypothesis that CAMAs are intracellular. Chlamydiae are known to co-occur with other bacteria in *Acanthamoeba* cells (*39*), although whether *Simkania* and *Endozoicomonas* share the same *P. acuta* cells is unclear. The close spatial proximity between them suggests the different bacterial species are interacting within the coral holobiont. Interestingly, both *Endozoicomonas* and chlamydiae are fish pathogens, forming cysts in the gills and skin and negatively affecting breathing (*40*). Such cysts strongly resemble CAMAs, although there is no record of *Endozoicomonas* and chlamydiae infecting the same fish hosts.

### Horizontal (*Endozoicomonas*) and vertical (*Kistimonas*) transmission of CAMA bacteria

The presence of CAMAs and of all four *Endozoicomonas* ASVs in both the F1 and F2 generations shows that this symbiotic association persists across generations and may be vertically transmitted to asexually produced larvae. However, we were not able to observe any CAMAs in freshly released F2 larvae (Figure S7). Because we may miss CAMAs when only looking at sections, we further conducted 16S rRNA gene metabarcoding on whole larvae of the F1_6 genotype to assess the presence of *Endozoicomonas* (Table S2). Of the 179 ASVs detected in these samples, none were assigned to the *Endozoicomonas* genus or to the Endozoicomonadaceae family (Table S4), suggesting the larvae do not contain CAMAs or *Endozoicomonas*. This contrasts with previous data showing the presence of CAMAs and of *Endozoicomonas* in larvae of distinct *P. acuta* genotypes sampled from Orpheus Island (*28*).

To further investigate the vertical transmission in the genotypes sampled by Damjanovic et al. (*28*) from Orpheus Island, we processed stored larval samples from that study belonging to the two genotypes OI2 and OI3 (Figure S2B, Table S1). FISH confirmed the presence of CAMAs in the larvae (Figure 4A-B), which also hybridized to the *Endozoicomonas* FISH probe (Figure 4C-D). We sampled CAMAs from both genotypes using LCM (Figures S1 and S4B) and examined their bacterial composition with 16S rRNA gene metabarcoding (Tables S2 and S3B). In the OI2 genotype, two ASVs assigned to *Kistimonas* made up more than 80% of the reads (Figure 4E). Only one nucleotide differs between the two ASVs, suggesting they may be a single *Kistimonas* strain. These two ASVs were also present in the OI3 genotype at a lower relative abundance (Figure 4E). Other ASVs in these samples may be contaminants that were too abundant to be removed bioinformatically as initial biomass was low in these samples. *Kistimonas* belongs to the Endozoicomonadaceae family, is phylogenetically close to *Endozoicomonas*, and has previously been isolated from marine invertebrates (*41*). Intriguingly, *Kistimonas* were not detected by Damjanovic et al. (2020), although the larvae were collected at the same time and from the same parent colonies. Therefore, we re-analyzed data from Damjanovic et al (*28*), comprised of whole larvae, recruits and adults, using an updated SILVA database (v138 vs v132). Among the seven *Endozoicomonas* ASVs previously reported, three were assigned to *Kistimonas* in our new analysis (Figure 4F), two of which perfectly matched the two ASVs found in our excised CAMA samples (ASVs 09 and 10). This could be due to the relatively recent description of *Kistimonas*, which might not have been included in v132 of the SILVA database. Importantly, only *Kistimonas*, and no *Endozoicomonas*, were detected in whole larvae and recruits, while both genera are present in the later life stages (Figure 4F).

**Figure 4:**
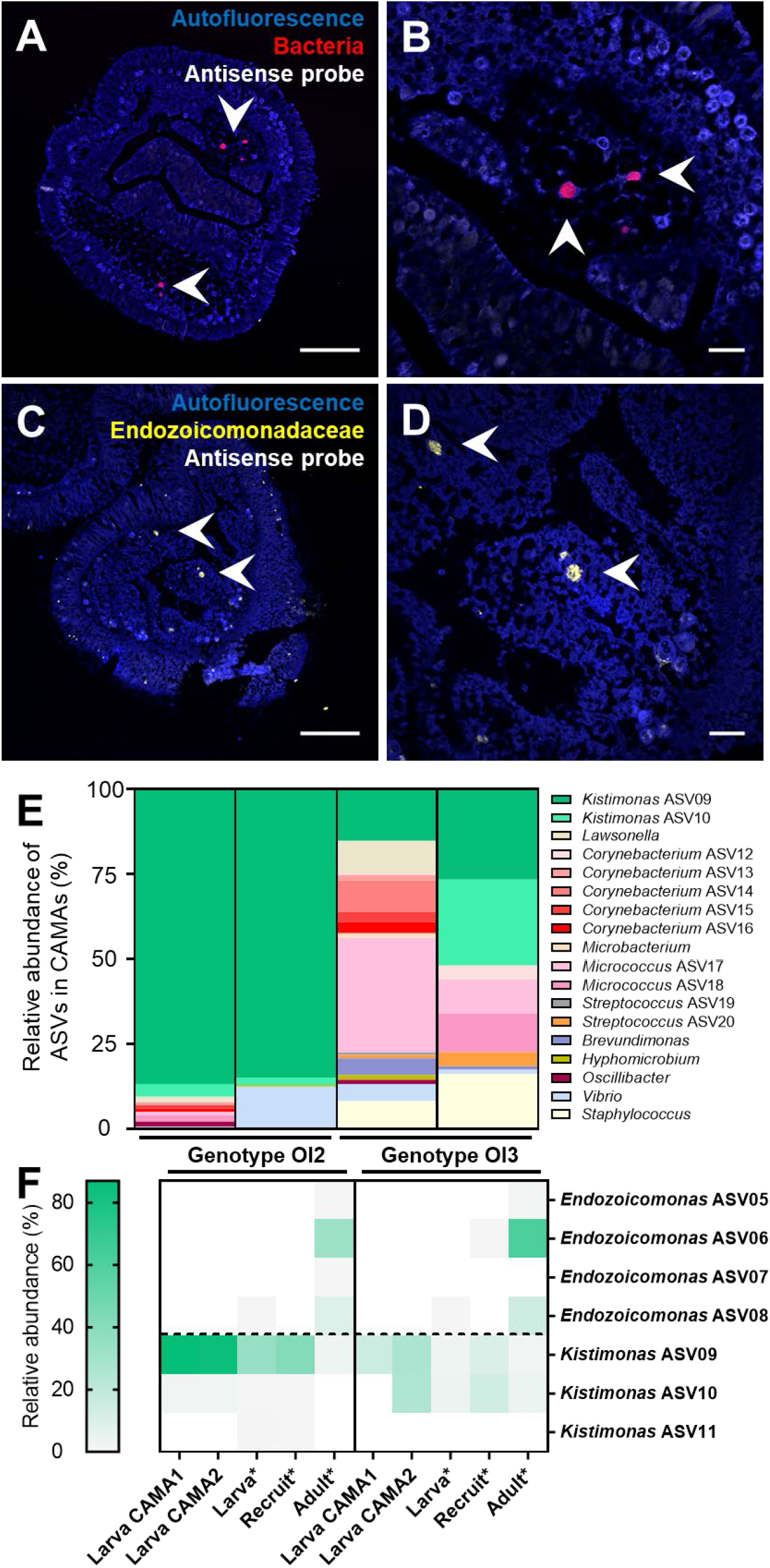
*Kistimonas* is the major CAMA taxon in larvae of the OI2 and OI3 genotype and is vertically transmitted. A-D: CAMA location by FISH on sectioned larvae. Arrowheads point at CAMAs. Blue: autofluorescence; yellow: End663 probe (Endozoicomonadaceae); red: EUB338-mix probe (all bacteria). Scale bars: 100 μm for A, C; 20 μm for B, D. E: Relative abundance of bacterial ASVs in CAMAs of larvae (OI2 and OI3 genotypes) isolated by LCM. Each bar is a single sample. F: Relative abundance of *Kistimonas* and *Endozoicomonas* ASVs in CAMAs, whole larvae, recruits, and adults of the OI2 and OI3 genotypes. Columns with an asterisk (Larva, Recruit, Adult) contain data re-analyzed from Damjanovic et al. (*28*).

Taken together, these data show that *Kistimonas* are vertically transmitted in genotypes OI2 and OI3, while *Endozoicomonas* are acquired horizontally in genotypes F1_6, OI2 and OI3, likely at the recruit or adult stage. Whether the vertical transmission of *Kistimonas* involves CAMAs or individual bacteria able to establish a new population remains unknown.

Interestingly, the *Simkania* ASV detected in the F2 adult samples of the F1_6 genotype was detected in every whole larval sample of the same genotype, with relative abundances ranging from 0.2 to 0.9% (Table S4). This suggests that *Simkania* is efficiently transmitted to asexually produced larvae in the F1_6 genotype, and that CAMAs containing *Simkania* may have been missed when sampling the F1 adult samples. *Endozoicomonas* is likely to have an environmental life stage, while *Kistimonas* and *Simkania* may be obligate symbionts. This is expected for *Simkania*, as all described chlamydiae depend on a eukaryotic host for propagation, whereas *Kistimonas*, just like *Endozoicomonas*, has been cultured *in vitro* from several marine hosts (*41*). These different transmission modes are likely to have impacted genome evolution and host-microbe interactions.

### CAMA bacteria belong to undescribed *Endozoicomonas* and *Simkania* species

To assemble the genomes and investigate the functional potential of uncultured CAMA bacteria, metabarcoding samples were used for shotgun sequencing and metagenomic analyses. Insufficient material was available of the OI2 and OI3 genotype larval samples to conduct metagenomics, thus only F1_6 samples were processed. Three metagenome-assembled genomes (MAGs) were recovered, one from the F1 generation sample, and two from the F2 generation samples. MAG data are detailed in Table 1. One MAG from each generation was assigned to *Endozoicomonas* (Pac_F1 and Pac_F2a), while the second MAG from the F2 generation sample was assigned to *Simkania* (Pac_F2b), consistently with our metabarcoding results. Pac_F1 and Pac_F2a genome sizes are similar with other *Endozoicomonas* genomes (*10, 25, 26*), while Pac_F2b is significantly smaller, which is consistent its classification as a member of the intracellular chlamydiae (*42*). Pac_F2b’s genome size is like other animal-associated chlamydiae (1-1.5 Mb), whereas protist-associated chlamydiae usually possess larger genomes (2-3 Mb) (*42, 43*). This may be due to the higher stability of a multicellular organism and relaxed selection pressure on more bacterial genes leading to deeper genomic erosion (*44*).

**Table 1:** Summary of the three metagenome-assembled genomes recovered in CAMAs sampled from adults of the F1_6 genotype.

| MAG | Pac_F1 | Pac_F2a | Pac_F2b |
| --- | --- | --- | --- |
| Genus assigned by GTDB-Tk | *Endozoicomonas* | *Endozoicomonas* | *Simkania* |
| Size (bp) | 6,938,003 | 5,907,264 | 1,247,175 |
| Coverage (mean value) | 2,063.5x | 3,539x | 500.7x |
| Completeness (%) | 94.25 | 88.76 | 88.02 |
| Contamination (%) | 1.65 | 2.48 | 0.21 |
| G+C content (%) | 49 | 51.3 | 43.9 |
| N50 | 6,009 | 6,281 | 41,614 |
| L50 | 314 | 248 | 12 |
| Number of contigs | 1,770 | 1,443 | 47 |
| Number of coding sequences (CDS) | 5,947 | 5,256 | 1,074 |
| Number of rRNAs | 10 | 5 | 1 |
| Number of tRNAs | 41 | 50 | 34 |

Because *Endozoicomonas* are so widespread in marine invertebrates, and particularly corals (*24*), we assessed their taxonomic placement. A full length 16S rRNA gene (1,536 bp) was recovered from Pac_F2a and two non-overlapping partial sequences were retrieved from Pac_F1 (401 bp and 1,129 bp respectively). ASV01 and ASV04 had 100% identity with Pac_F1 (1,129 bp fragment) and Pac_F2a, respectively. A phylogenetic tree based on more than 1,000 Endozoicomonadaceae 16S rRNA gene sequences shows our two strains cluster together with other *Endozoicomonas* strains isolated from *P. damicornis* (*45*) and *S. pistillata* (*16, 21*) corals (Figure S8). This indicates a certain degree of host specialization in the Pocilloporidae family (*21*). However, this contrasts with other *animal-Endozoicomonas* associations, with host phylogenies rarely matching symbiont phylogenies (*21, 25, 26*), which is further evidence that *Endozoicomonas* possess an environmental life stage, are not obligate symbionts, and are not undergoing genome streamlining. Pac_F1 and Pac_F2a form a well-supported clade together with *E. acroporae* (isolated from *Acropora* sp. corals), and to a lesser extent *E. atrinae*, and *E. elysicola*, although they likely represent a new species (~95-97% identity with *E. acroporae, E. atrinae*, and *E. elysicola*). A phylogenetic tree based on 120 bacterial gene markers from 15 *Endozoicomonas* genomes confirmed this trend (Figure 5, Table S5A). Two clades were observed, with coral-associated *Endozoicomonas* present in both. Pac_F1 and Pac_F2a clustered together and were closest to two *Endozoicomonas* genomes recently sequenced from *S. pistillata* CAMAs (*21*). Average nucleotide identity (ANI) and average amino acid identity (AAI) showed strong similarity between Pac_F1 and Pac_F2a (98% and 96%, respectively) (Figure S9A-B). The highest values were obtained with *Stylophora-* (ANI 83%, AAI 78%) and *Acropora-*associated (ANI 80%, AAI 70%) *Endozoicomonas*. Overall, this data suggests that Pac_F1 and Pac_F2a are two different strains that belong to a novel, undescribed *Endozoicomonas* species, which is closely related to other Pocilloporidae-associated *Endozoicomonas*.

**Figure 5:**
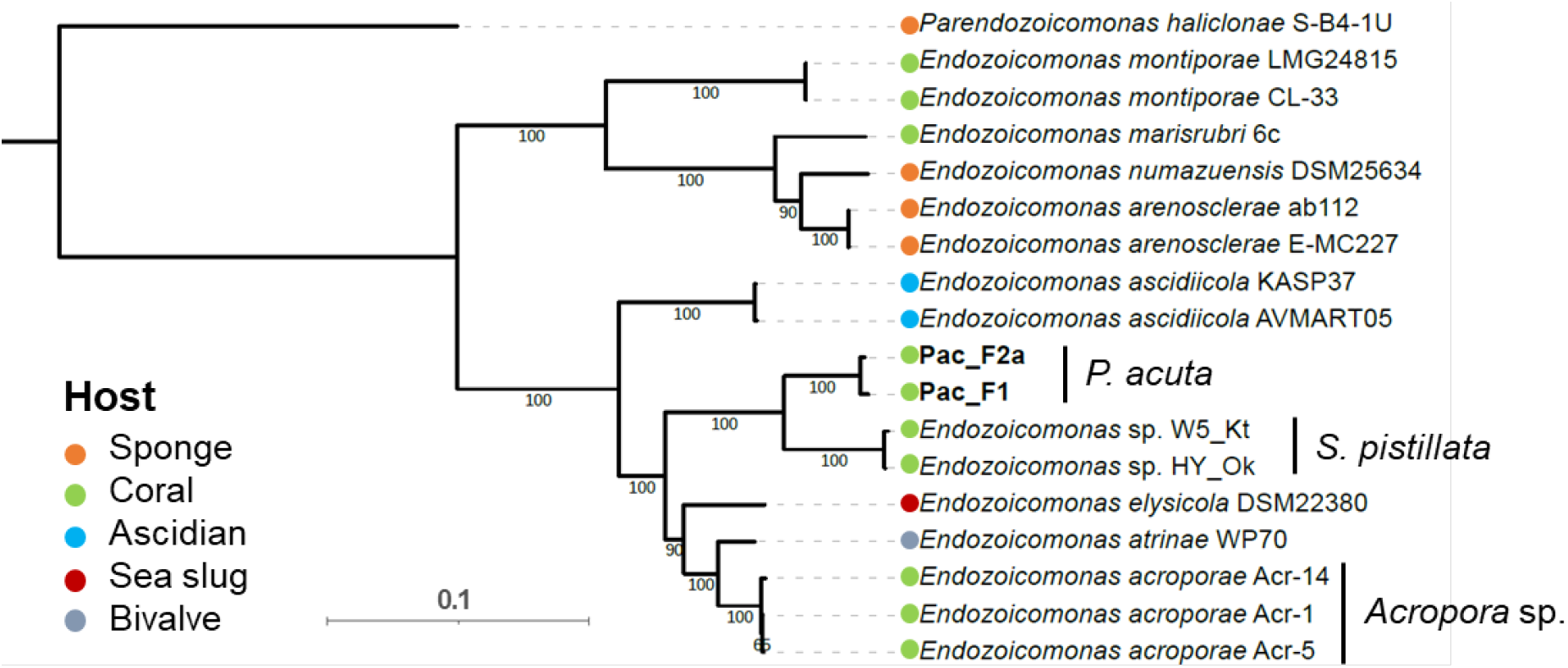
Two MAGs sequenced from CAMAs belong to new species *Endozoicomonas*. Maximum-likelihood phylogeny of *Endozoicomonas* based on 120 marker genes, 15 *Endozoicomonas* genomes, and one *Parendozoicomonas* genome (outgroup). Bootstrap support values based on 1000 replications are provided. Additional data on the reference genomes is available in Table S5A.

Only a partial 16S rRNA sequence (662 bp) was retrieved for Pac_F2b, so we created a phylogenetic tree based on 15 conserved marker genes, along with other chlamydiae and Simkaniaceae genomes (Table S5B). As suggested by the GTDB-tk classification, Pac_F2b falls in the same clade as *Simkania negevensis* and is closest to a MAG isolated from the coral *Cyphastrea* sp. sampled in subtropical Lord Howe Island (Australia) (Figure 6 and S10). AAI was highest with the MAG isolated from *Cyphastrea* sp. (75%) and *S. negevensis* (64%) (Figure S11). ANI values were too low (<75%) to be reliable. This suggests that Pac_F2b in an undescribed species that belongs to the *Simkania* genus.

**Figure 6:**
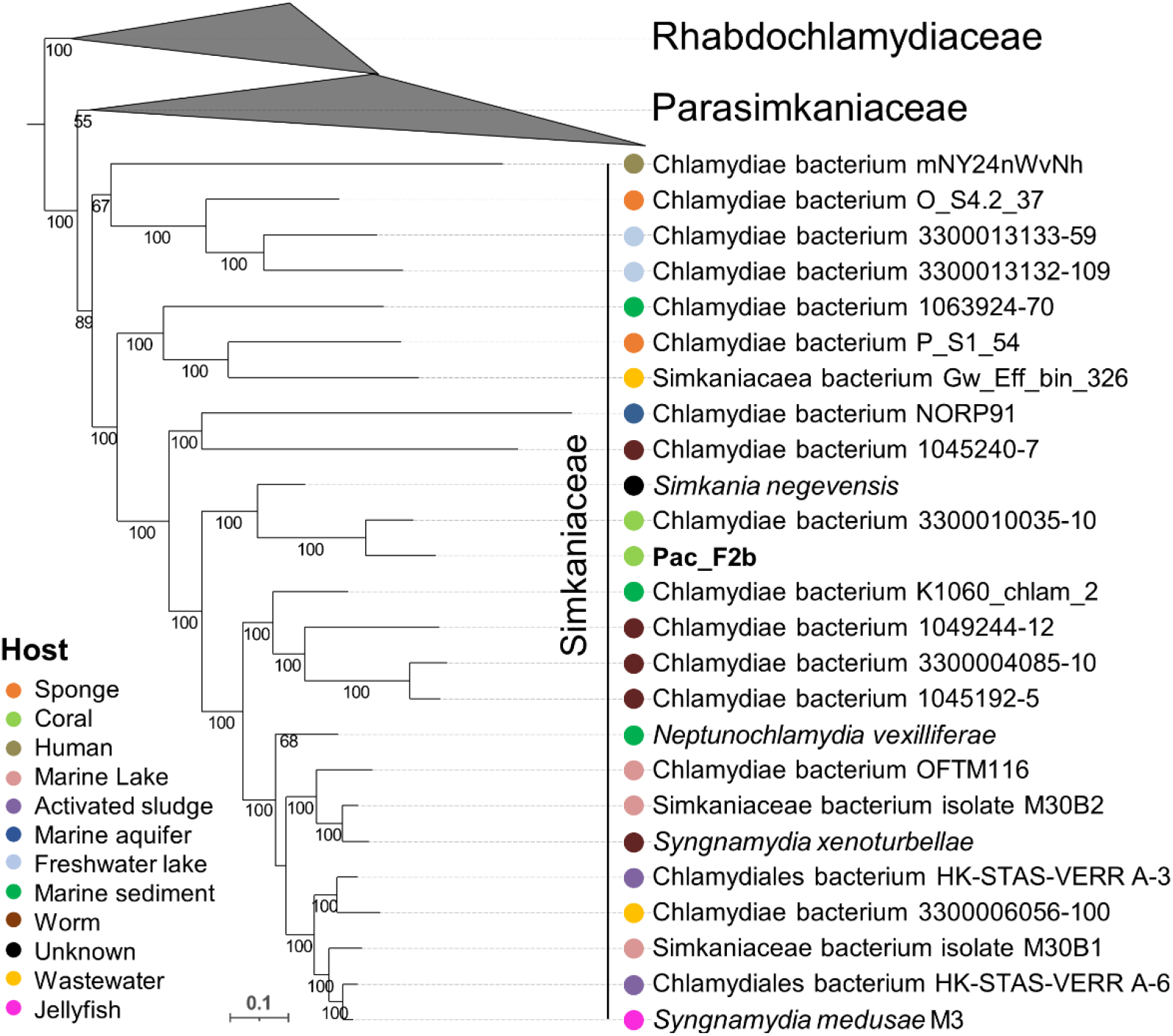
One MAG sequenced from CAMAs belongs to a new species of *Simkania*. Chlamydial maximum-likelihood phylogeny based on 15 conserved non-supervised orthologous groups (NOGs) in 24 Simkaniaceae, 8 Parasimkaniaceae, and 21 Rhabdochlamydiaceae (outgroup) genomes. Bootstrap support values based on 1000 replications are provided. Additional data on the reference genomes is available in Table S5B. A tree containing the entire chlamydial phylogeny is available in Figure S10.

### *Endozoicomonas* may provide vitamins to their coral host and protect it against pathogens

The functional potential of the three MAGs recovered from CAMAs of *P. acuta* was investigated through KEGG, RAST and InterProScan analyses and is summarized in Figure 7. The completeness of selected KEGG pathways is provided in Figure S12. Detailed Prokka and eggNOG-mapper annotations are provided in Dataset S1. We first focused on genes for host-symbiont interactions in the two *Endozoicomonas* MAGs, Pac_F1 and Pac_F2a (Figure 7A). First, a total of nine secondary metabolites were bioinformatically predicted across the two MAGs (Table S6). Only one showed similarity with a known biosynthetic gene cluster, which encodes the antibacterial protein rhizomide (*46*). The predicted secondary metabolites were also annotated by Prokka and were identified as antimicrobial proteins, such as gramicidin, tyrocidine, and microcin. Additionally, several near-complete secretion systems were detected, including a type II, type III, and type VI (T2SS, T3SS, and T6SS, respectively). Note that a T6SS was only detected in Pac_F1. All three secretion systems may be involved in host infection and/or host cell entry, although their effectors remain unknown. While T2SSs and T3SSs are routinely observed in *Endozoicomonas* (*10, 21, 25*), this is only the second report of a T6SS in a coral-associated *Endozoicomonas* (*21*). These secretion systems, along with the secondary metabolites, may play a role in host colonization, outcompeting other bacteria present in the coral holobiont, as well as preventing other bacteria from invading CAMAs and promoting the maintenance of these structures. High numbers of genes encoding eukaryotic-like proteins were also detected (Table S7), including more than 100 ankyrin-repeat proteins in each MAG. These are also found in other *Endozoicomonas* (*10, 21, 25*), are often effectors of secretion systems, and hypothesized to facilitate host-bacteria interactions by mediating bacterial protein-eukaryotic host protein interactions (*47*). Finally, both MAGs possess the complete machinery to synthesize type IV pili (T4P). While never discussed in earlier publications, the pathway is also complete in other *Endozoicomonas* genomes (Table S8). However, the gene encoding the actual pilus, *pilA*, is missing in *Parendozoicomonas haliclonae*. T4P are involved in bacterial motility and may play a role in host colonization, but they also support the formation of biofilms and aggregates (*48*). Specifically, *pilY1* encodes an adhesin that may be involved in host recognition. Therefore, we hypothesize that T4P are involved in the formation and maintenance of *Endozoicomonas* CAMAs. As *E. acroporae* forms CAMA-like structures in culture when incubated with dimethylsulfoniopropionate (DMSP) (*10*), it is possible that metabolic signals, such as DMSP, regulate the *in hospite* expression of T4P synthesis genes and in turn the formation of CAMAs.

**Figure 7:**
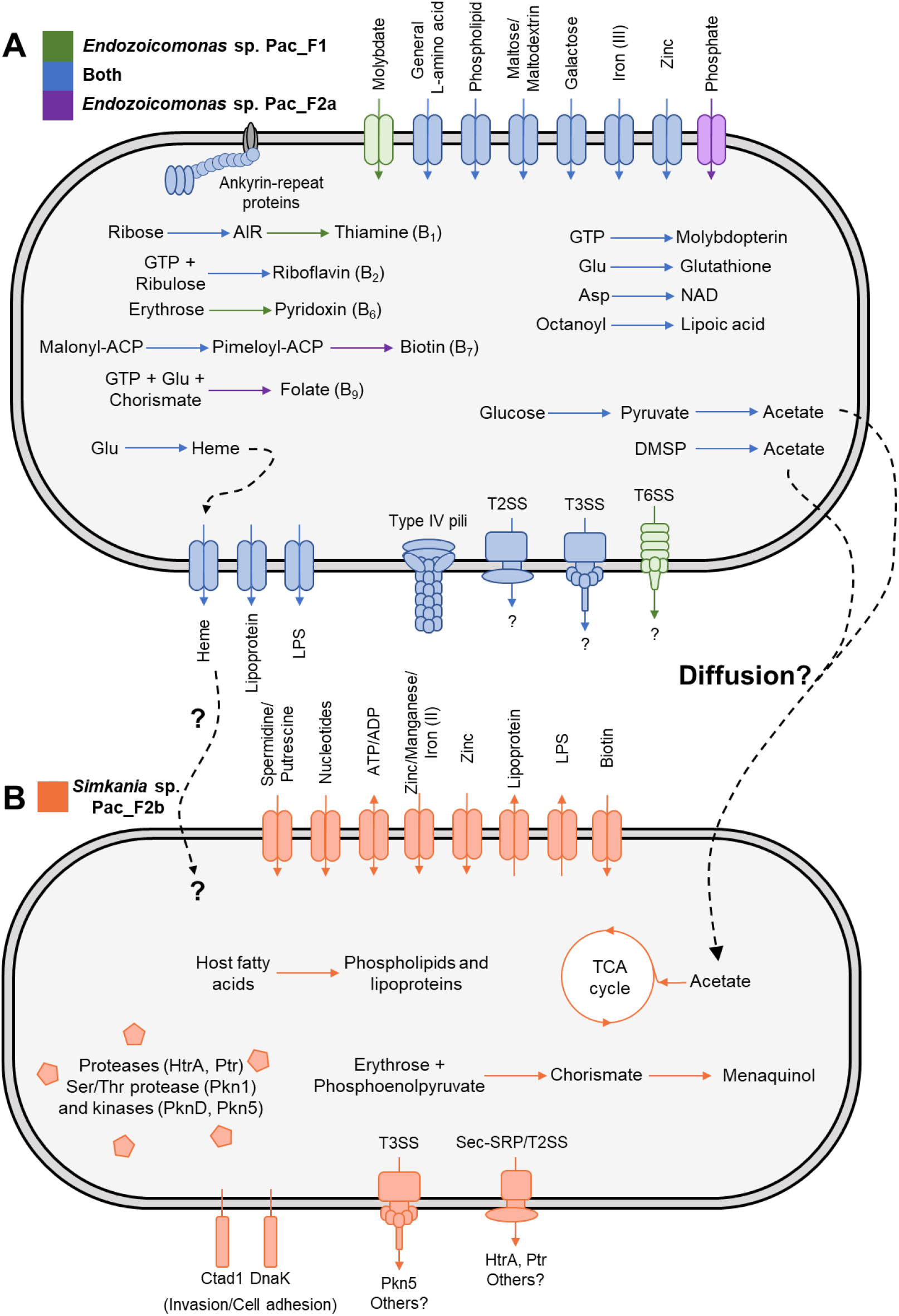
Overview of the genomic potential of the three MAGs recovered from *P. acuta* CAMAs, *Endozoicomonas* sp. Pac_F1 (A) and Pac_F2a (A), and *Simkania* sp. Pac_F2b (B). The pathways represented here are either complete or only lack one gene (Figure S12). Dashed arrows represent hypothetical interactions between *Endozoicomonas* and *Simkania* (Figure S13). Sec-SRP: Sec translocase and signal recognition particle pathway; T2SS: type II secretion system; T3SS: type III secretion system; T6SS: type VI secretion system; TCA: tricarboxylic acid; AIR: aminoimidazole ribotide; DMSP: dimethylsulfonioproprionate; LPS: lipopolysaccharide; NAD: nicotinamide adenine dinucleotide.

Both *Endozoicomonas* MAGs encode pathways for the synthesis of several B vitamins, including thiamine (B1), riboflavin (B2), pyridoxin (B6), biotin (B7) and folate (B9). Both corals and Symbiodiniaceae cannot produce B vitamins and are dependent on external sources. B vitamin provisioning from bacteria has been shown in hematophagous insects (*49*) and marine algae (*50*), and B vitamin synthesis pathways have been detected in other *Endozoicomonas* (*21, 25, 26*). Further, thiamine and pyridoxin synthesis was upregulated in *E. marisrubri* incubated with coral host extracts (*25*). It is therefore likely that *Endozoicomonas* provide B vitamins to its coral host and/or Symbiodiniaceae, although how these vitamins are exported remains elusive. As in other *Endozoicomonas* (*21, 26*), we identified a wide range of ABC transporters, notably for zinc, iron, amino acids, maltose, galactose, or heme. *Endozoicomonas* MAGs recently obtained from CAMAs in *S. pistillata* showed the presence of genes involved in phosphate transport, metabolism, and storage, as well as a phosphotransferase system, which led to the hypothesis that CAMAs may play a role in phosphate cycling and balance in the coral holobiont (*21*). Here, we only found genes encoding an ABC transporter for phosphate and a cellobiose phosphotransferase system, but no other genes that may suggest a role in phosphate metabolism. Finally, we also detected several pathways for the synthesis of molecules involved in antioxidant processes, including lipoic acid, heme (incorporated into hemoproteins, such as catalases and peroxidases), and glutathione. Antioxidant synthesis is a trait of high interest in the design of bacterial probiotics for the mitigation of coral bleaching (*51, 52*), as the overproduction of reactive oxygen species by Symbiodiniaceae is thought to be the main driver of coral bleaching (*53*). Thus, *Endozoicomonas* is likely to be a beneficial symbiont for its host, potentially providing vitamins, antioxidants, and antimicrobial molecules. As the relative abundance of *Endozoicomonas* often diminishes during thermal stress (*54*), the loss of its associated benefits may exacerbate the negative effects of marine heatwaves, including the increase in abundance of opportunistic pathogens.

### *Simkania* may benefit from *Endozoicomonas* for growth and survival

The *Simkania* MAG Pac_F2b exhibited reduced metabolic abilities (Figure 6B), which is consistent with other chlamydiae and their intracellular lifestyle (*42*). It possesses all the genes for glycolysis, the tricarboxylic acid (TCA) cycle, and the pentose phosphate pathway. Additionally, Pac_F2b encodes the genes for the transformation of acetate into acetyl-CoA, potentially enabling this *Simkania* strain to use acetate as a substrate for the TCA cycle and as a carbon source (Figure S12). In a similar way, sponge-associated chlamydiae can use acetoin as an energy source (*55*). Interestingly, both *Endozoicomonas* MAGs can produce acetate via two pathways (Figure S13): (i) through the conversion of acetyl-coA, produced by pyruvate oxidation, into acetate; (ii) through the conversion of DMSP into methylmercaptopropionate by DmdA, and subsequently into acetate, although not all the genes in this pathway were found. Because acetate can freely diffuse across membranes, we hypothesize that excess acetate produced by *Endozoicomonas* present in neighboring CAMAs can be acquired by *Simkania* for its own growth. Like most chlamydiae, Pac_F2b lacks the genes for *de novo* nucleotide biosynthesis and can only synthesize few amino acids (alanine, aspartate, and glutamate), but possesses a complete shikimate pathway. Other hallmark chlamydial genes were detected, including a T3SS, adhesins, ATP/ADP translocases, and virulence effectors such as proteases and kinases (Figure 7B). An orthogroup analysis revealed that 518 of Pac_F2b’s genes (almost 50%) were present in more than 90% of all publicly available chlamydial genomes and 204 genes were shared by two other *Simkania* genomes (Dataset S2). This suggests that Pac_F2b’s overall genomic content is similar to that of other chlamydiae. The only vitamin/cofactor Pac_F2b can produce is menaquinol, which is common in chlamydiae (*55*) and plays a role in bacterial growth in *C. trachomatis* (*58*). Pac_F2b may import other vitamins from host cells or adjacent *Endozoicomonas* through vitamin transporters such as BioY, present in Pac_F2b. Unlike most other chlamydiae (*55*), it does not encode the heme biosynthesis pathway. Although the genome sequence of the Pac_F2b MAG is not complete, it is highly unlikely that the full heme pathway is present. None of the heme biosynthesis genes could be detected in the closest relative, the *Cyphastrea-associated* Chlamydiae bacterium 3300010035-10, either and these genes are scattered at six loci throughout the chromosome in *Simkania negevensis*. Heme is essential for many cellular processes, including antioxidant functions, as well as infectivity and virulence in many pathogens. For example, heme production is essential for the intracellular survival of *Brucella abortus* (*59*). Heme-deficient strains of *Staphylococcus aureus* show reduced tissue colonization in mice compared to wild-type strains (*60*), although they persist more easily intracellularly, perhaps because of lower toxin production (*61*). The specific role of heme in chlamydiae remains unknown, although mutations in the *hemG* gene, involved in heme synthesis, were associated with increased infectivity in *C. trachomatis* (*62*). In *P. acuta, Simkania* may acquire heme from the host, or from neighboring *Endozoicomonas*, which possess the complete heme biosynthesis pathway and an ABC transporter for heme. However, no heme import system was detected in Pac_F2b and how heme would cross CAMA or *Simkania’*s membranes remains unclear. Alternatively, it is possible that the lack of heme synthesis in Pac_F2b makes this strain less virulent and able to persist in inclusions in *P. acuta*, similarly to *S. aureus*.

In summary, it is possible that *Simkania* is reliant on *Endozoicomonas* for acetate, which *Simkania* can use to stimulate its growth, as well as heme and vitamins. Interactions between bacteria within a holobiont are known in insects where endosymbiotic bacteria share a single cell, *e.g. Buchnera/Serratia* in the aphid *Cinara cedri (63*) or *Tremblaya/Moranella* in the mealybug *Planococcus citri* (*64*) (note that in the latter, *Moranella* resides within *Tremblaya* cells). In both cases, the synthesis of some amino acids requires a patchwork of genes present in one symbiont or the other. The potential metabolic interactions we uncovered in *P. acuta* may explain the spatial proximity of the two types of CAMAs, and why we did not observe any *Simkania* CAMAs on their own. Nevertheless, the presence of *Simkania* without *Endozoicomonas* in larvae samples of the F1_6 genotype (Table S4) suggests that potential *Endozoicomonas*-*Simkania* interactions are not necessary for *Simkania* survival. However, *Simkania* was in low relative abundance in these samples (< 1%) and was not observed forming CAMAs (Figure S6). This suggests that either its growth is impeded, or that it is instead dependent on host-produced acetate and/or heme. In the latter hypothesis, *Simkania* would not be restricted to aggregate around *Endozoicomonas* and may be able to source host compounds in different coral cell types, hence the absence of CAMAs in these samples. Simkaniaceae and Endozoicomonadaceae co-occur in corals of the *Acropora, Stylophora* and *Pocillopora* genera (*21, 30, 35, 65*). These corals cover the complex and robust clades within the Scleractinia. The interactions suggested by our genomic data may thus be widespread in coral holobionts and should be studied further.

### Suitability for the development of probiotic treatments

We provided here a comprehensive overview of the biology of CAMAs in the coral *P. acuta*. CAMAs are located at the tip of the tentacles, contain members of the Endozoicomonadaceae and Simkaniaceae families, exhibit mixed-mode transmission, and genomic data suggests they bear a wide range of functions. This further establishes *Endozoicomonas* as a widespread mutualistic symbiont of corals and we provide additional evidence that *Endozoicomonas* symbionts vary in their genetic potential and life strategies. This is also the first evidence, both spatial and genomic, for CAMAs constituted by different microbes suggesting a relevance for interactions between CAMAs for CAMA and host functioning.

Our study provides crucial insights into the biology of these endosymbionts, which may be used for the development of probiotics for the mitigation of coral bleaching. Specifically, several aspects suggest the strong suitability of *Endozoicomonas*. First, they may have several beneficial functions, including an antagonistic effect towards other bacteria, and the production of B vitamins and antioxidants. Second, our data strongly suggests that *Endozoicomonas* is intracellular, and shows efficient spreading from colony to colony, as all individuals from two successive generations were infected. This stability is essential if probiotics are to lastingly colonize coral hosts and successfully spread through coral populations, without having to repeatedly inoculate probiotics (*4, 51*). However, as they are not located in the gastrodermis in *P. acuta*, it is unclear whether any beneficial function would affect Symbiodiniaceae (*e.g.*, through the transfer of antioxidants to the gastrodermis) and thus be relevant for coral bleaching. *Kistimonas*, through its vertical transmission, would also be a candidate of choice and needs to be investigated further. Future work should focus on culturing these strains*. Kistimonas* may be more challenging to culture if vertical transmission has resulted in some genome degeneration and higher host dependence than *Endozoicomonas*. Cultured isolates will allow us to perform targeted analyses to confirm their functional potential, as well as reinoculation experiments to trial their potential efficiency as probiotics.

## Material and methods

### Coral husbandry and aquarium conditions

For the F1_6, C2_12 and R2_8 genotypes, 3 parent colonies were collected from 3 reefs in July 2017, Feather reef (colony ID: 202B F1_6; Lat: −17.51854 Long: 146.38940), Coates reef (colony ID: 307A C2_12; Lat: −17.18850 Long: 146.37180) and Rib reef (colony ID: 211D R2_8; Lat: −18.47122 Long: 146.87360), along the central Great Barrier Reef in Australia (Table S1, Figure S2A). The colonies were brought back to the National Sea Simulator at the Australian Institute of Marine Science (AIMS) in Townsville, QLD, Australia. Parent colonies were kept in aquaria, running an annual temperature profile based on historical SST records (1985-2012) for the mid-shelf Central GBR obtained from National Oceanic and Atmospheric Administration (NOAA) Coral Reef Watch (CRW) daily global 5km (0.05 degree) satellite product v3.1 (Figure S14). The annual temperature profile targeted an annual average temperature for the central GBR of 26.14 ± 0.23°C (± 1 daily SD); an annual range from 23.16 to 29.88°C, with a summer heat stress accumulation equivalent to 0.95°C-weeks (NOAA CRW v3.1). Corals were maintained in a semi-recirculating system with a daily turnover of ~300%, and circulation within tanks was achieved with two pumps (Maxspect 350 series) generating gyres at opposite ends of the tanks; partial CO_2_ pressure was maintained at a daily average value 464 ± 40.94 ppm (± 1 daily SD) allowing for diurnal variability driven by natural photosynthesis and respiration cycles; daily light cycles were set to match local conditions with daily and seasonal variations in solar intensity and lunar cycles to maintain synchronicity of planulation patterns for *P. Acuta* colonies. From July to October 2017, fully developed F1 larvae, released asexually by the parent colonies, were collected with flowthrough collection devices on the outflow of isolation tanks and settled separately under constant temperature-controlled flow-through seawater onto pre-conditioned aragonite settlement plugs. From September 2018 to February 2019, F2 larvae, generated asexually by the F1 colonies (12-16 months old at that point), were collected with flow-through collection devices on the outflow of isolation tanks and settled separately under constant temperature-controlled flow-through seawater onto pre-conditioned aragonite settlement plugs. Throughout their life, both generations were kept in the same mesocosm systems under the same annual conditions as parent colonies.

For the OI2 and OI3 genotypes, parent colonies were collected from Orpheus Island, in the central Great Barrier Reef in Australia (Table S1, Figure S2B), and maintained and sampled as previously described (*28*).

### Sampling

For adult colonies, small coral fragments were snapped off the colonies with forceps, rinsed with 0.22 μm FSW and placed in cryovials. Forceps were rinsed in 80% ethanol between sampling each colony. Two to three fragments per colony were placed in one cryovial. Three colonies were sampled per genotype. For FISH and LCM, branches were fixed for 24 hrs in 4% paraformaldehyde (PFA) in 0.22 μm-filtered seawater (FSW), rinsed twice in FSW, and stored in 50% ethanol-PBS at −20°C. For transmission electron microscopy (TEM), branches were fixed for 24 hrs in 4% PFA + 0.1% glutaraldehyde in FSW, rinsed once in FSW, once in PBS 1X, and stored in PBS 1X at 4°C. Following fixation, coral branches were decalcified in EDTA 10%. EDTA was renewed every two days, and samples were kept at 4°C on a rotating wheel, until there was no skeleton left (around two weeks). Samples were then rinsed in PBS 1X, and stored at 4°C in PBS 1X.

Larvae were sampled as previously described (*28*). Briefly, before planulation, colonies were maintained in individual acrylic aquaria that received indirect natural sunlight and coarsely filtered seawater. A filter was fitted at each outlet of the acrylic tank to collect released planulae. Each larva was washed with FSW transferred from the filter into an Eppendorf tube (FISH) or cryovial (metabarcoding) using a sterile pipette tip and as much water as possible was removed without disturbing the larvae. For FISH and LCM, larvae were fixed for 24 hrs in 4% PFA in FSW, rinsed twice in FSW, and stored in 50% ethanol-PBS at −20°C. For 16S rRNA gene metabarcoding, larvae were snap frozen and stored at −80°C. For FISH, around 10 larvae per genotype were fixed. For metabarcoding, larvae were pooled (three per tube), immediately snap-frozen following sampling and kept at −80°C.

### FISH on slides

Sample processing, sectioning of larvae and decalcification of branches were performed by the Melbourne Histology Platform (University of Melbourne). First, samples were dehydrated in 50% ethanol for 1 hr, 70% ethanol for 1 hr, 90% ethanol for 45 min, 100% ethanol for 1 hr (three times), 50% ethanol and 50% xylene for 45 min, xylene for 45 min (twice), and paraffin for 50 min (three times). Samples were subsequently embedded in solid paraffin, and 3-μm sections were cut using a Microm HM 325 rotary microtome and plain microscopic slides (Menzel, Germany). Each slide had four serial sections. FISH was then performed as previously described (*28*), except final probe concentration during hybridization was 5 ng/μL. See Table S9 for probe sequences and formamide concentrations (*16, 38, 66, 67*). Slides were mounted in CitiFluor™ CFM3 mounting medium (proSciTech, Australia) containing 3 μg/uL 4’,6-diamidino-2-phenylindole dihydrochloride (DAPI; Merck, Germany) to stain nucleic acids, covered with a coverslip and sealed with clear nail polish. Slides were kept at 4°C until observation.

### Whole-mount FISH

Single polyps were dissected in PBS 1X using Dumont tweezers under a dissecting microscope. Tissue was cleared of autofluorescence by incubating polyps in 50% methanol for 10 min, 75% methanol for 10 min, 90% methanol for 10 min, 100% methanol for 10 min, 90% methanol + 0.1% triton for 10 min, 75% methanol + 0.1% triton for 10 min, 50% methanol + 0.1% triton for 10 min, PBS 1X + 0.1% triton for 10 min, and stored at 4°C in PBS 1X.

The whole-mount FISH protocol was adapted from previously published protocols (*28, 68*). Samples were first permeabilized in 70% acetic acid for 2 min and twice rinsed in PBS 1X for 5 min. Samples were then incubated in PBS 1X + 0.1% triton for 10 min, PBS 1X for 5 min, pepsin at 0.2 mg/mL in HCl 0.01 M at 37°C for 30 min, PBS 1X for 5 min, HCl 0.2 M for 12 min, and Tris-HCl 20 mM for 10 min. Hybridization was performed in the dark for 3 hrs at 46°C in 2 mL of hybridization buffer (0.9 M NaCl, 20 mM Tris-HCl pH 7.2, 0.01% SDS, 5 ng/mL of probe; see Table S9 for formamide concentration). A negative control with no probe was included. Samples were then washed for 15 min at 48°C in washing buffer (20 mM Tris-HCl pH 7.2, 0.01% SDS; see Table S9 for NaCl concentration). Samples were then rinsed twice in ice-cold milliQ water and deposited onto 8-well coverslip-bottom slides (ibidi, USA). A drop of milliQ water was deposited onto each polyp to avoid drying. Slides were kept at 4°C until observation.

### Confocal laser scanning microscopy

Slides were observed on a Nikon A1R confocal laser scanning microscope (Nikon, Japan) with the NIS325 Element software. Virtual band mode was used to acquire variable emission bandwidth to tailor acquisition for specific fluorophores. The fluorophores Cy3 and Atto550 were excited using the 561 nm laser line, Atto647 using the 640 nm laser line, DAPI using the 405 nm laser line, and the coral autofluorescence using the 488 nm laser line with a detection range of 570-620 nm for Cy3 and Atto550, 660-710 nm for Atto647, 500-550 nm for coral autofluorescence and 420-480 nm for DAPI. For three-dimensional reconstructions of Z-stacks (Figures 1A and 3B), sections were acquired using Z steps of 3 μM with the 10X objective and 1.1 μM with the 20X objective. Nd2 files were processed using ImageJ. Linear adjustments of brightness and contrast were performed when necessary and applied to the entire image and to each channel independently. Channels were then merged together. Z-stacks were projected in two-dimensional images using the ‘Max Intensity’ projection type.

### Transmission electron microscopy

Following decalcification, single polyps were dissected in PBS 1X using Dumont tweezers under a dissecting microscope. Samples were washed in distilled water 3 x 10 min followed by post-fixation with 1% osmium tetroxide in ddH_2_O for 1 hr. Samples were then dehydrated in an acetone series of 10, 20, 40, 60, 80 and 100% acetone. Two further exchanges of 100% acetone were undertaken before infiltrating and embedding the samples in Spurr’s resin. Thin (100 nm) sections were cut on a Leica UC7 ultramicrotome and collected on copper slot grids. The sections were post-stained with 1% uranyl acetate in water for 10 min and 3% lead citrate for 2 min, and imaged on an FEI Tecnai Spirit transmission electron microscopy equipped with an Eagle CCD camera (Thermofisher). CAMAs were only detected in one polyp of the C2_12 genotype.

### Laser capture microdissection of CAMAs

Genotypes F1_6 (adults, generations F1 and F2) and O2 (larvae) were chosen to perform laser capture microdissection (LCM) to extract CAMAs. For each group, three replicates containing either one branch or three larvae were processed. For each replicate, 10 slides each containing 8 sections were processed (~80 sections per replicate in total). FISH was performed as described above, using the EUBmix-338-atto550 probe. Slides were not mounted and left without coverslip. To check that FISH had worked and to take photos comparing before and after LCM, one slide per group was covered with a coverslip (without any mounting medium) and sealed with clear nail polish for CLSM imaging as described above. The nail polish was then carefully dissolved by applying small amounts of acetone with a cotton swab and the coverslip was removed. All slides were processed by LCM the next day to avoid any loss of signal. LCM was performed on a PALM Laser Microdissector (Zeiss, Germany), using the PALM Robo 3.2 software, and a 20X air objective. CAMAs were detected using a TRITC filter (excitation 550 ± 12.5 nm; emission 605 ± 35 nm). Individual CAMAs were captured using a UV laser (355 nm) into the caps of 200 μL AdhesiveCap Clear tubes (Zeiss, Germany). These caps are filled with clear adhesive material that ensures sample retention. CAMAs from all slides from one replicate were all collected in one tube. For each replicate, tissue areas without CAMAs were also separately captured as a control. DNA extraction was then conducted within the cap, using the Arcturus® PicoPure® DNA Extraction Kit (Applied Biosystems, USA). 155 μL of Reconstitution Buffer into one vial of Proteinase K to dissolve the enzyme. 10 μL of the Proteinase K solution was then added to each cap and incubated at 65°C for 18 hrs. Tubes were then centrifuged for 5 min at 10,000 × *g* to bring samples down into the tubes and stored at −20°C. Two to three caps containing no sample, but that were open in the LCM facility to capture air contamination, were also included as extraction blanks.

### DNA extractions of whole larvae

Six replicates each containing 3 larvae were processed. DNA extractions were performed using a salting-out method as previously described (*69*). Three blank DNA extractions were conducted as negative controls. A mock community, ZymoBIOMICS Microbial Community DNA Standard (Zymo Research), was included to check sequencing and processing quality.

### 16S rRNA gene metabarcoding

Hypervariable regions V5-V6 of the 16S rRNA genes were amplified using the primer set 784F (5’ <u>GTGACCTATGAACTCAGGAGTC</u>AGGATTAGATACCCTGGTA 3’) and 1061R (5’ <u>CTGAGACTTGCACATCGCAGCC</u>RRCACGAGCTGACGAC 3’). Adapters were attached to the primers and are shown as underlined. Bacterial 16S rRNA genes were PCR-amplified on a SimpliAmp Thermal Cycler (Applied Biosystems, ThermoFisher Scientific). Each reaction contained 1 μL of DNA template, 1.5 μL of forward primer (10 μM stock), 1.5 μL of reverse primer (10 μM stock), 7.5 μL of 2x QIAGEN Multiplex PCR Master Mix (Qiagen, Germany) and 3.5 μL of nuclease-free water (Thermofisher), with a total volume of 15 μL per reaction. Three triplicate PCRs were conducted for each sample and three no template PCRs were conducted as negative controls. PCR conditions for the 16S rRNA genes were as follows: initial denaturation at 95°C for 3 min, then 18 cycles of: denaturation at 95°C for 15 s, annealing at 55°C for 30 s, and extension at 72°C for 30 s; with a final extension at 72°C for 7 minutes. Samples were then held at 4°C. Following PCR, triplicates were pooled, resulting in 45 μL per sample. Metabarcoding library preparation was conducted as previously described (*70*) and sequencing was performed at the Walter and Eliza Hall Institute (WEHI) in Melbourne, Australia on one MiSeq V3 system (Illumina) with 2×300bp paired-end reads.

### Bacterial 16S rRNA gene analysis

QIIME2 v 2020.11 (*71*) was used for processing 16S rRNA gene sequences. The plugin demux (*71*) was used to create an interactive plot to visualize the data and assess the quality, for demultiplexing and quality filtering of raw sequences. The plugin cutadapt (*72*) was used to remove the primers and MiSeq adapters. Plugin DADA2 (*73*) was used for denoising and chimera checking, trimming, dereplication, generation of a feature table, joining of paired-end reads, and correcting sequencing errors and removing low quality reads (Q-score < 30). Summary statistics were obtained using the feature-table to ensure processing was successful. Taxonomy was assigned by training a naive Bayes classifier with the feature-classifier plugin (*71*), based on a 99% similarity to the V5-V6 region of the 16S rRNA gene in the SILVA 138 database to match the 784F/1061R primer pair used (*74*). Mitochondria and chloroplast reads were filtered out. Analyses and graphs were performed using Rstudio version 2022.02.2 and the phyloseq package (*75*). Metadata file, taxonomy table, phylogenetic tree and ASV table were imported into R to create a phyloseq object. Contaminant ASVs, arising from kit reagents and sample manipulation, were identified using the package decontam (*76*). The function ‘isNotContaminant’ was used as it is more stringent and more adequate for low-biomass samples. One replicate for each the Adult F1 condition, larvae OI2 and larvae OI3 were removed because of heavy contamination. It is worth noting that bacteria belonging to the *Brachybacterium* genus accounted for more than 90% of the contamination (Table S2), which were contaminants of the Qiagen PCR kit.

### Genome amplification of LCM samples and shotgun sequencing

To reach sufficient quantities for shotgun sequencing, DNA samples obtained through LCM needed to be amplified. Whole-genome amplification was carried out using a SeqPlex DNA Amplification Kit (Sigma-Aldrich, USA), which is specifically designed for low-quantity, fragmented samples. The same DNA samples that were used for 16S rRNA gene metabarcoding were used and, for each sample, 1.5 μL of each of the three replicates were pooled before amplification. This was only carried out using the F1_6 genotype samples (both generations F1 and F2), as there was not enough material left from the O2 and O3 genotype samples. Amplification was performed according to the manufacturer’s instructions, using 29 cycles in an end-point PCR. Intermediate DNA concentrations were measured using a Qubit Fluorometer and a Qubit high-sensitivity dsDNA quantification assay. Primer removal was also carried out using the SeqPlex DNA Amplification kit. Final DNA quality and quantities were checked on a TapeStation (Agilent, USA). 16S rRNA gene metabarcoding of the amplified samples showed similar results to the non-amplified samples, confirming that amplification bias was minimal (Table S3A).

Samples were then sent to the Ramaciotti Centre for Genomics (Sydney, Australia) for sequencing. Library preparation was performed with a NextFLEX Rapid DNA Seq 2.0 Prep kit and sequenced on a NovaSeq 6000 SP 2×100 bp Flowcell Illumina platform.

### Data analysis of metagenomics

Each sample was run on two different lanes, so reads from each lane were first merged, according to their direction (i.e., Forward reads from both lanes were merged, and Reverse reads from both lanes were merged). Adapters were removed from the raw data using Cutadapt v4.0 (*72*) with default settings. Quality control was performed using FastQC v0.11.9 (*77*) and low-quality sequences (phred score < 30) were trimmed using Trimmomatic v0.36 (*78*) (LEADING:30 HEADCROP:10 MINLEN:70). High-quality reads were mapped against a *Pocillopora acuta* draft genome (*79*) using Bowtie2 v2.4.5 with default parameters (*80*) to remove host related reads from the metagenome. Mapped reads were then removed using Samtools v1.11 (*81*). Only paired-end host-removed reads were used for metagenome assembly. Metagenome assembly was carried out using MEGAHIT v1.2.9 (*82*) with a minimum contig length of 1000 bp and the following k-mers: 21, 33, 55, 77. Trimmed reads were then mapped back to the assembly using Samtools v1.11 (*81*). Assembled contigs were binned using the binning (with metabat2, concoct and maxbin2 tools) and bin_refinement (with >70% completeness and <10% contamination as cut-off parameters) modules of MetaWRAP v1.3.2 (*85*). Bin quality and taxonomy were assessed using CheckM2 v0.1.3 (*86*) and GTDB-Tk v2.1.0 (*87*), respectively. These bins were reassembled to further improve the contiguity and bin completeness and contamination stats using the reassemble_bins module SPAdes implemented in MetaWRAP v1.3.2. The final taxonomy of individual contigs on a per bin level was assessed using CAT/BAT v5.2.3 (*84*), and any contig belonging to a different phylum than the taxonomy assigned by GTDB-Tk was manually removed. Bin coverage was obtained using CoverM v0.6.1 (https://github.com/wwood/CoverM) using the “genome” option.

### Genome annotation

Gene prediction and annotation of bins was performed using Prokka v1.14.6 with default settings (*88*), RAST v2.0 (*89*), KEGG-mapper Reconstruct (*90*), eggNOG-mapper v2.1.6 with a cut-off e-value of 1.10^-5^ (*91*) with the eggNOG database v. 5.0 (*92*), and InterProScan v5.55 with Pfam domain annotations (*93*). For Pac_F2b (*Simkania* MAG), Prokka annotation was performed using the “--protein” option with a well-annotated set of chlamydial genes as initial reference. Secondary metabolites were predicted using antiSMASH v6.1.1 (*94*). For the annotation of DMSP demethylases (DmdABC), conserved domains were double-checked with web-based CD-search using e-value cut-off 1e-5 (*95*). Average Nucleotide Identities (ANI) and Average Amino acid Identities (AAI) of all three MAGs were calculated using a genome-based matrix calculator (*96*).

### Phylogenetic analyses

16S rRNA gene sequences were recovered from MAGs using barrnap as implemented in Prokka. The 16S rRNA gene phylogenetic tree of the Endozoicomonadaceae family was constructed using full-length 16S rRNA gene sequences from the SILVA v132 database. For Pac_F2a, a full-length 16S rRNA gene (1536 bp) was recovered and used for the phylogenetic tree construction. For Pac_F1, two fragments were recovered, of 1129 bp and 401 bp respectively and only the 1129-bp fragment was used. A MAFFT alignment was created using Geneious Prime v2019.1.3. This alignment was used to generate a maximum likelihood phylogenetic tree with 1000 ultrafast bootstraps using IQ-Tree v2.2.0.3 (*97*) with the best model TIM3e+R6, selected by ModelFinder wrapped in IQ-tree (*98*). Sequences belonging to the *Zookishella* genus were chosen as an outgroup. For the *Endozoicomonas* whole-genome phylogenetic tree, GTDB-Tk v2.1.0 (*87*) was used to create an alignment based on 120 gene markers. Nucleotide sequences were chosen because all the genomes belong to the same genus. The genome of *P. haliclonae* was selected as an outgroup. The reference genomes are listed in Table S5. A maximum likelihood phylogenetic tree was constructed based on the alignment with IQ-Tree v2.1.2 (*97*) using the best model LG+F+R3, selected by ModelFinder wrapped in IQ-tree (*98*), and 1000 ultrafast bootstrap replicates (*99*).

For comparative genome analysis of the Pac_F2b MAG, a dataset of high quality chlamydial genomes based on Dharamshi et al. (*55*) was used. This dataset was complemented by additional chlamydial genomes available on NCBI (last accessed on 01/09/2022). Only genomes with a completeness >70% (determined with CheckM2 v0.1.3 (*86*)) and average nucleotide identities <95% (determined with FastANI v1.33 (*100*)) were considered for the final dataset which included 139 chlamydial genomes (Table S5B). For further comparative analysis of the chlamydial genomes, all encoded protein sequences were clustered into orthologous groups (OGs) with OrthoFinder v2.5.2 with default parameters (*101*). In order to obtain the phylogenetic affiliation of the Pac_F2b MAG, a set of concatenated protein sequences from 15 non-supervised orthologous groups (NOGs) was used (*55*) (Table S10). It has been shown previously that the usage of these 15 NOGs retrieves the same topology for chlamydial phylogeny as the application of larger protein sets (*55*). Thus, the dataset from Dharamshi et al. (*55*) including a large outgroup of other members of the Planctomycetes, Verrucomicobia, Chlamydiae (PVC) superphylum was expanded by the Pac_F2b MAG and the additional genomes from NCBI. Proteins of these genomes belonging to the 15 NOGs were aligned to the existing alignments from Dharamshi et al. (*55*) with MAFFT v7.490 “--add” (*102*). The resulting single protein alignments were subsequently trimmed with BMGE v1.12 (*103*) (entropy 0.6) and concatenated. Maximum likelihood phylogeny was inferred with IQ-TREE v2.1.2 (*97*) with 1000 ultrafast bootstrap replicates (*99*) and 1000 replicates of the SH-like approximate likelihood ratio test (*104*) under the LG+C60+F+R4 model.

## Acknowledgments

We thank the Biosciences Microscopy Unit (University of Melbourne) and the Biological Optical Microscopy Platform (University of Melbourne) for the use of their confocal microscopes and laser capture dissection, and particularly Dr Gabriela Segal, Dr Shane Cheung and Dr Sam Mills for their valuable assistance. We thank the Melbourne Histology Platform (University of Melbourne), Laura Leone, and Lisa Foster for their assistance with sample sectioning. We also thank the Melbourne Research Cloud and the Life Science Compute Cluster (LiSC; http://cube.univie.ac.at/lisc) for providing the high-performance computing instances needed for this work. We are grateful to Dr Sanjida Topa and Ms Isini Buthgamuwa for their efforts in trying to culture *Endozoicomonas*.

## Author contributions

Conceptualization: JM, LB, MJHvO; Formal analysis: JM, KT, AC, GP, MH; Funding acquisition: MJHvO, NC; Investigation: JM, AvdM, KD, CRG, SS, NC; Methodology: JM; Supervision: LB, MJHvO; Visualization: JM, KT, AC; Writing - Original Draft Preparation: JM, MJHvO. All authors have read and edited the manuscript.

## Data availability

Raw data are available under NCBI BioProject IDs PRJNA891910 (Adult CAMAs, MiSeq and NovaSeq raw data, MAGs), PRJNA891898 (Larva CAMAs, MiSeq raw data), PRJNA891892 (Whole larva microbiome, MiSeq raw data).

