## Supplementary materials for "*Endozoicomonas*-chlamydiae interactions in cell-associated microbial aggregates of the coral *Pocillopora acuta*"

**This PDF file includes:**

Figs. S1 to S14

Tables S2, S7, and S10

Legends for Tables S1, S3, S4, S5, S6, S8, and S9

Legends for Dataset S1 and S2

**Other Supplementary Materials for this manuscript include the following:**

Tables S1, S3, S4, S5, S6, S8, and S9

Dataset S1 to S2

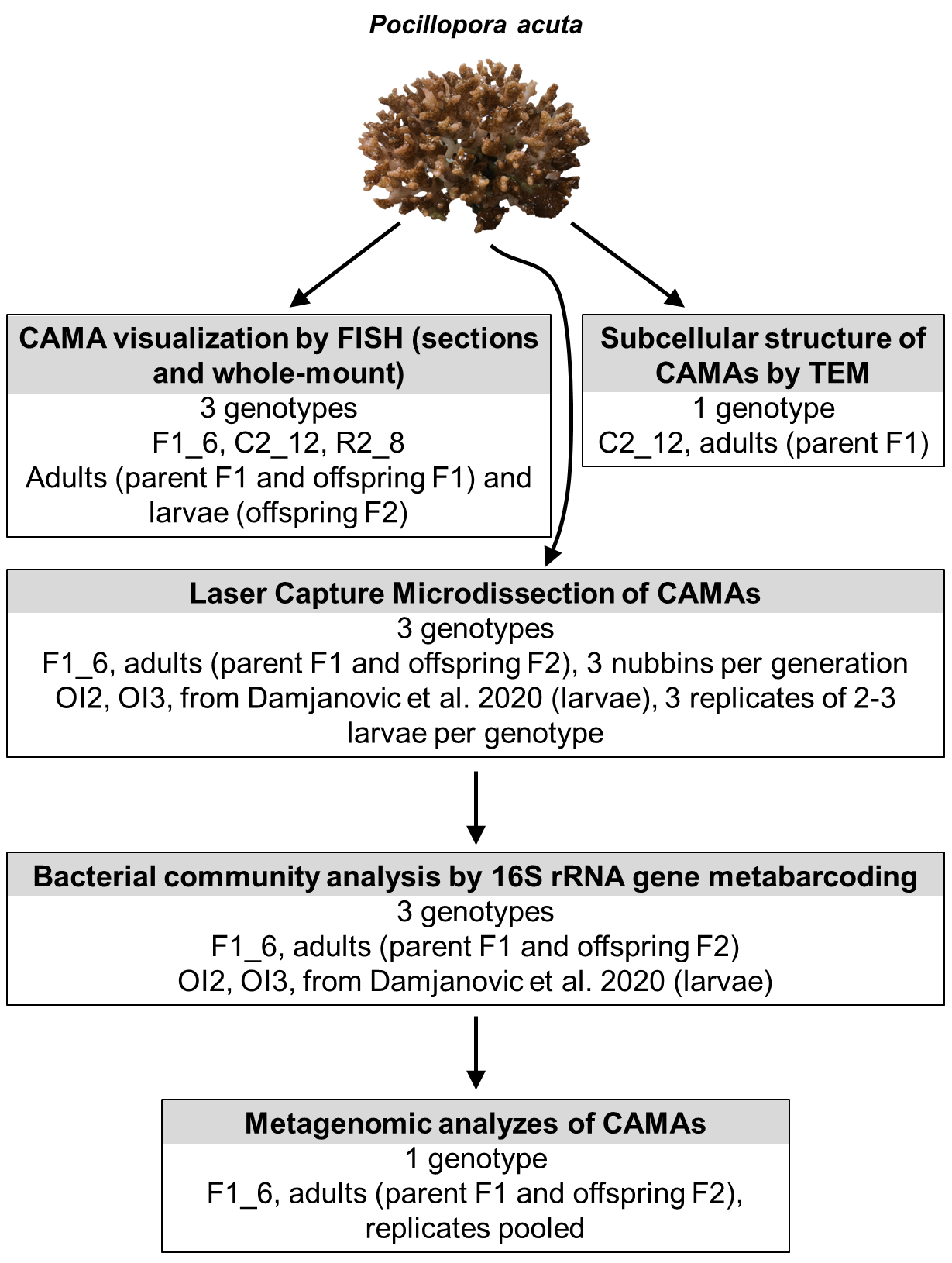

Fig. S1. Experimental design used in this study. Adults (F1 generation and F2 generation) and larvae (F2 generation) of three genotypes of *P. acuta* colonies bred in captivity (F1_6, C2_12, R2_8) were used for FISH. Adults (F1 generation) of the C2_12 genotype was used for TEM observations. Adults (F1 and F2 generation) of the F1_6 genotypes, as well as larvae sampled by Damjanovic et al. (2020) of two genotypes (OI2 and OI3) (*28)* were used to sample CAMAs by LCM. LCM samples were used for 16S rRNA gene metabarcoding for community profiling, and samples of the F1_6 genotype were further used for metagenomics analyzes. CAMA: Cell-associated Microbial Aggregates. FISH: Fluorescence *in situ* Hybridization; SEM: Scanning Electron Microscopy. LCM: Laser Capture Microdissection.

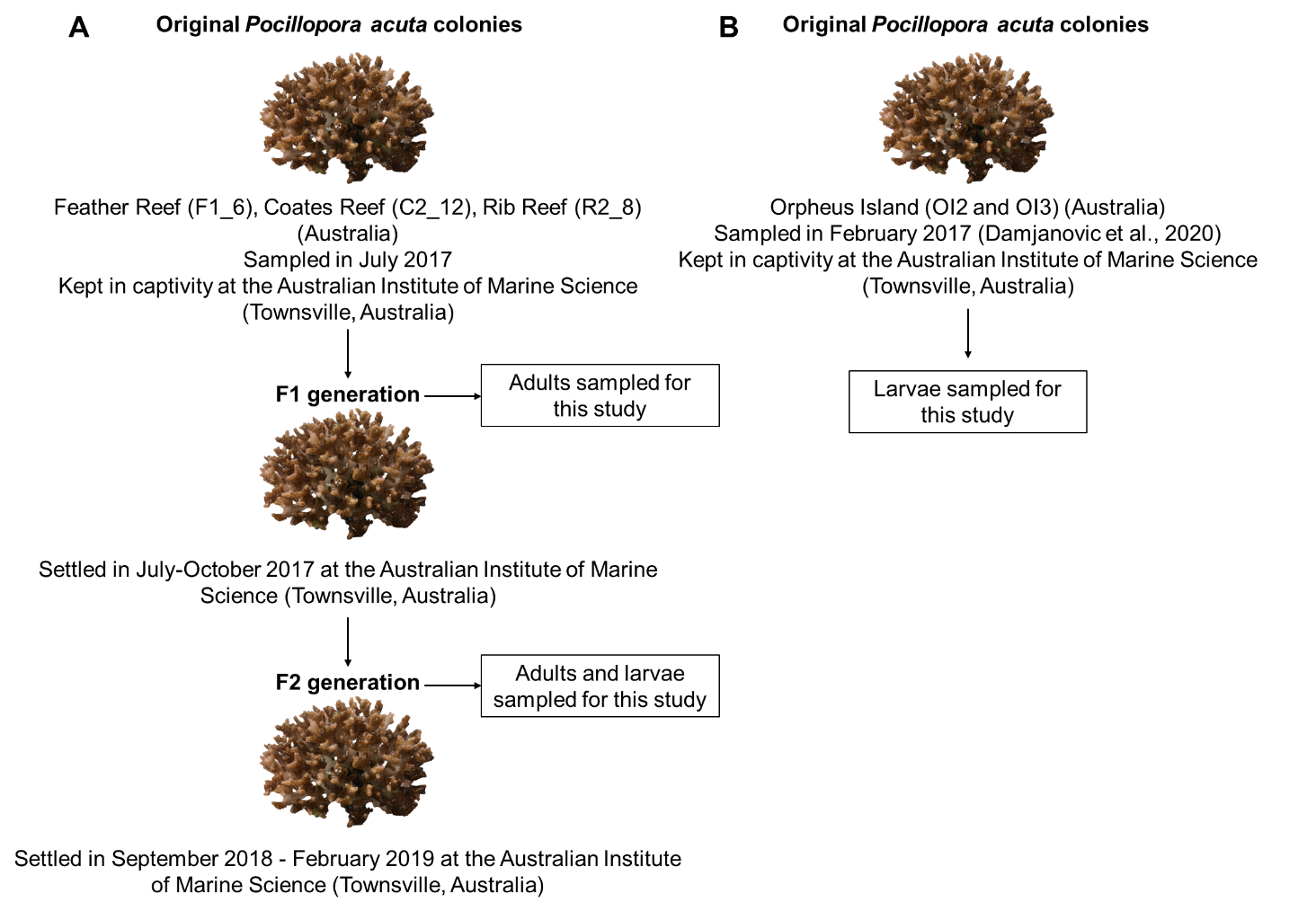

Fig. S2. Sampling performed for this study. A: Sampling of the F1_6, C2_12, and R2_8 genotypes, referred to in Figures 1, 2, 3, 5, 6. Following establishment of the original colonies in captivity, adults of the F1 and F2 generation, and larvae of the F2 generation (released by adults of the F2 generation) were sampled. B: Sampling of the OI2 and OI3 genotypes (performed by Damjanovic et al. (2020) (*28*)), referred to in Figure 4. Following establishment of the original colonies in captivity, larvae released by the adults were sampled. In the original study, the original adults and recruits were also sampled.

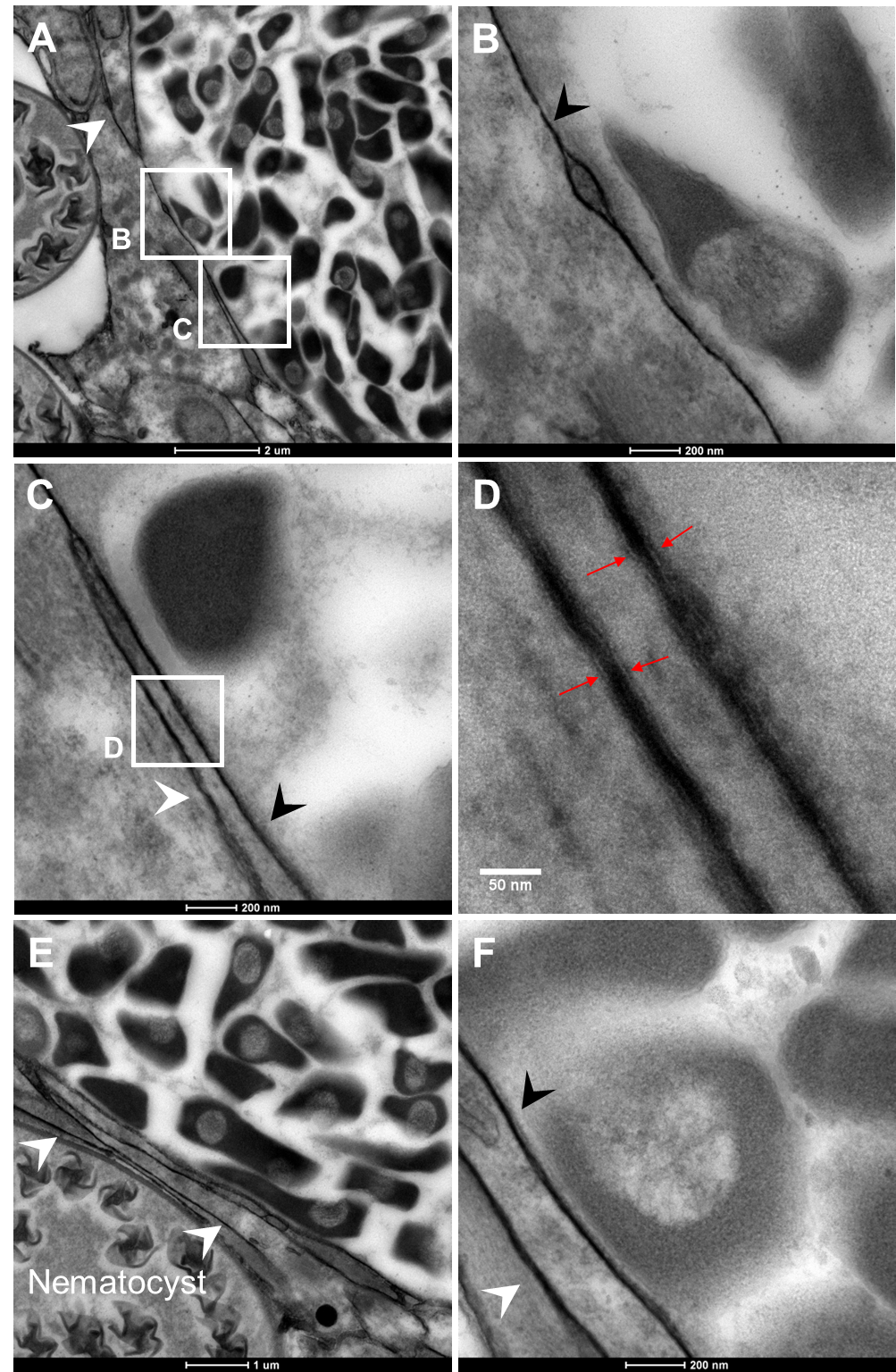

Fig. S3: Additional TEM photos of CAMAs. Black arrowheads point at a possible membrane surrounding the CAMAs. White arrowheads point at possible coral cell membranes. Red arrows point at lipid bilayers. B and C are magnifications of A. D is an enlargement of C.

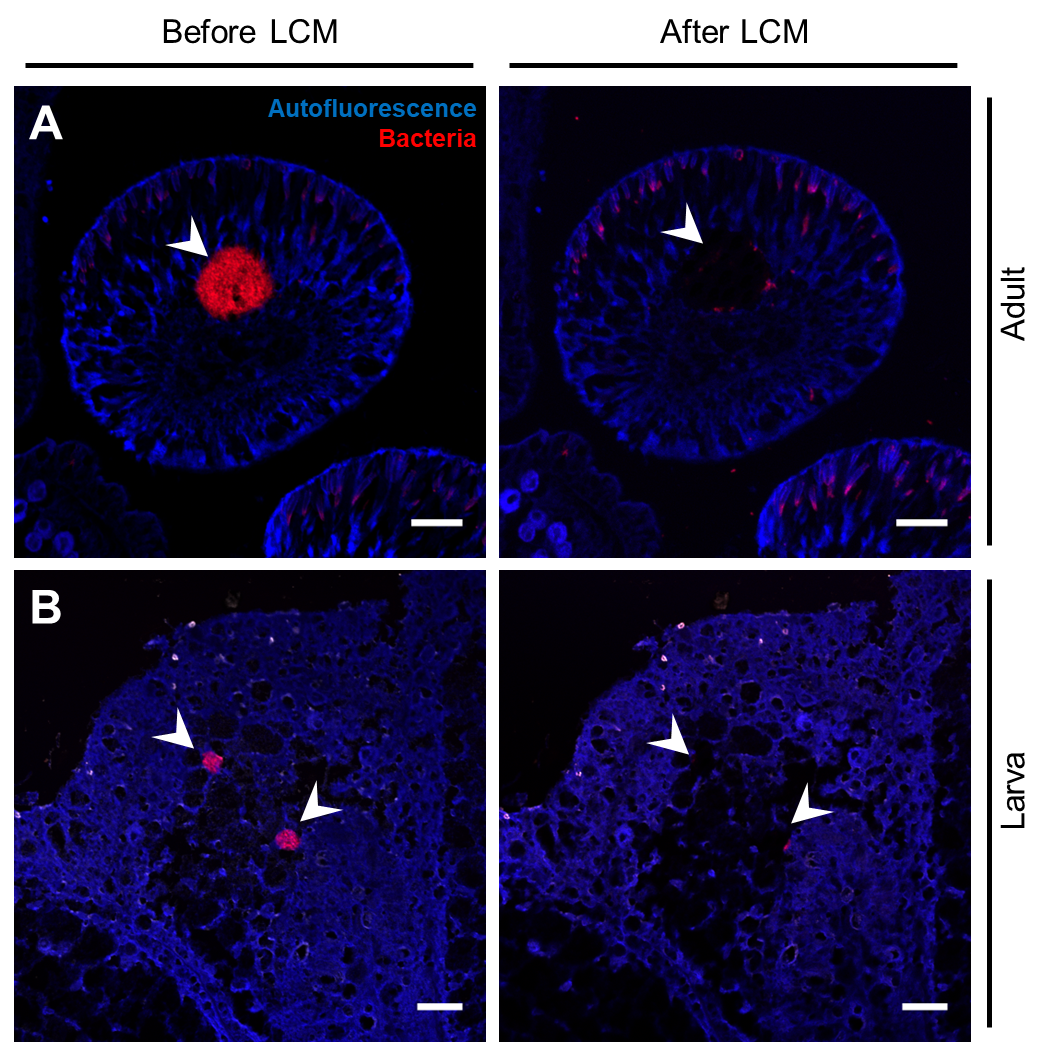

Fig. S4. Laser capture microdissection of CAMAs in adults (A, F1_6 genotype, F2 generation) and larvae (B, OI2 genotype). Arrowheads point at CAMAs (left panels) and captured CAMAs (right panels) in the same section. Blue: autofluorescence; red: EUB338-mix probe (all bacteria). Scale bars: 20 µm for A-D.

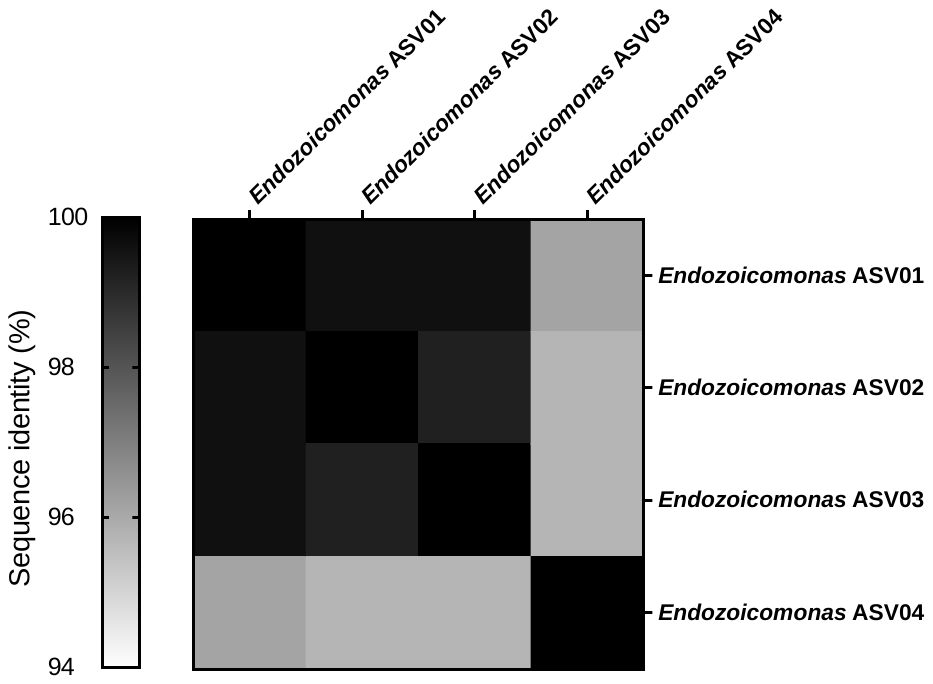

**Fig. S5:** Nucleotide identity between the four *Endozoicomonas* ASVs recovered in CAMAs sampled from F1_6 adult *P. acuta* corals.

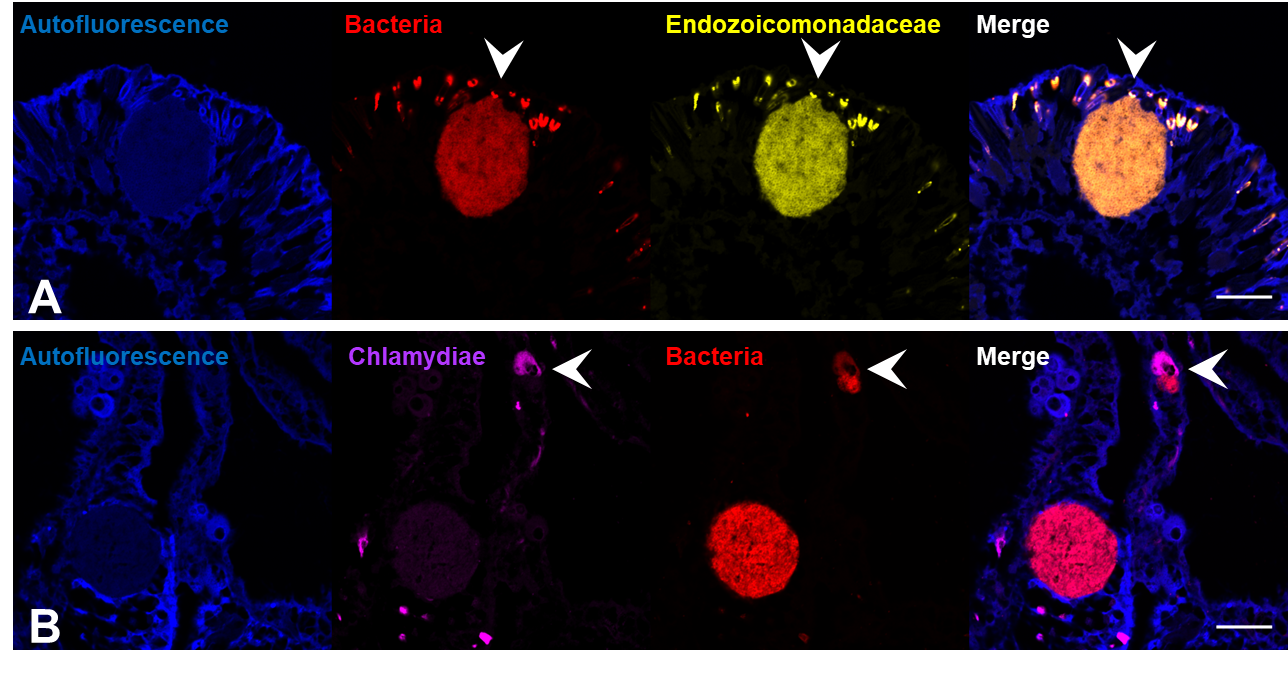

**Fig. S6:** Colocalization of bacterial and taxon-specific probes. All photos are from the F1_6 genotype. Generation: F1 (A), F2 (B). Blue: autofluorescence; yellow: End663 probe (Endozoicomonadaceae); magenta: Chls523 probe (chlamydiae), red: EUB338-mix (all bacteria). Scale bars: 20 µm for A, B. White arrowheads points at an Endozoicomonadaceae CAMA in A, and at a chlamydiae CAMA in B.

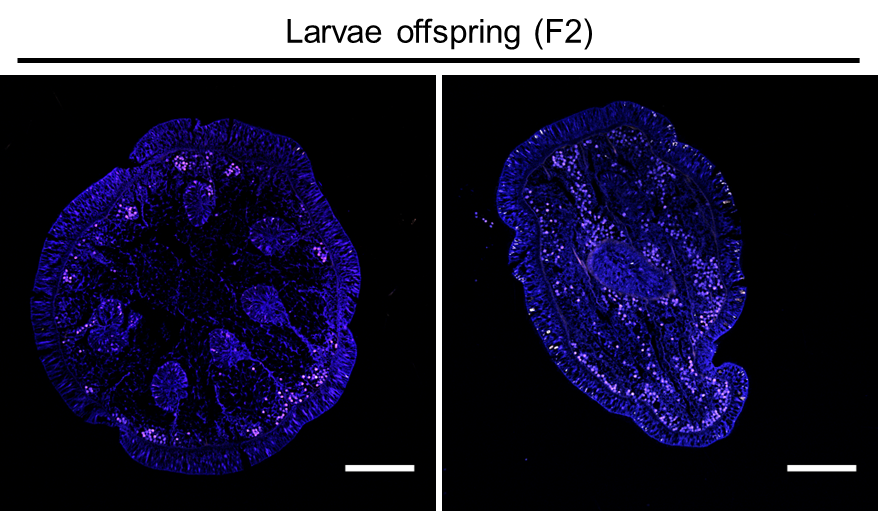

**Fig. S7:** Absence of CAMAs in larvae. FISH was performed on sectioned larvae (F2 generation). Genotype: C2_12 (left), F1_6 (right). Blue: autofluorescence; red: EUB338-mix probe (all bacteria); white: non-EUB probe (negative control). Scale bars: 200 µm.

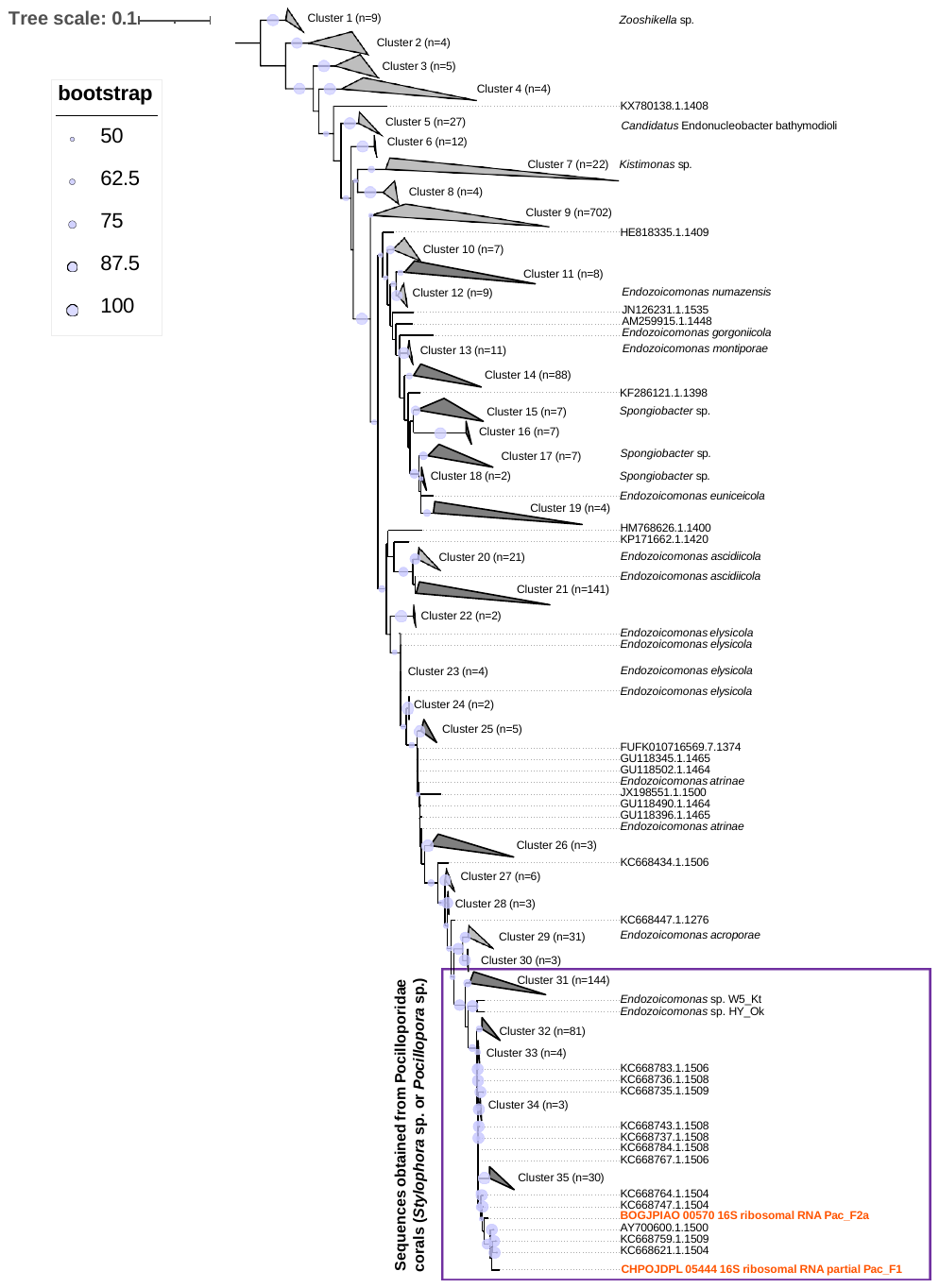

**Fig. S8:** Maximum likelihood phylogenetic tree showing the placement of Pac_F1 and Pac_F2a in the Endozoicomonadaceae family based on 1459 bacterial 16S rRNA sequences from the SILVA database, in addition to the two 16S rRNA sequences extracted from our two MAGs. Bootstraps values greater than 50% based on 1000 replications are provided.

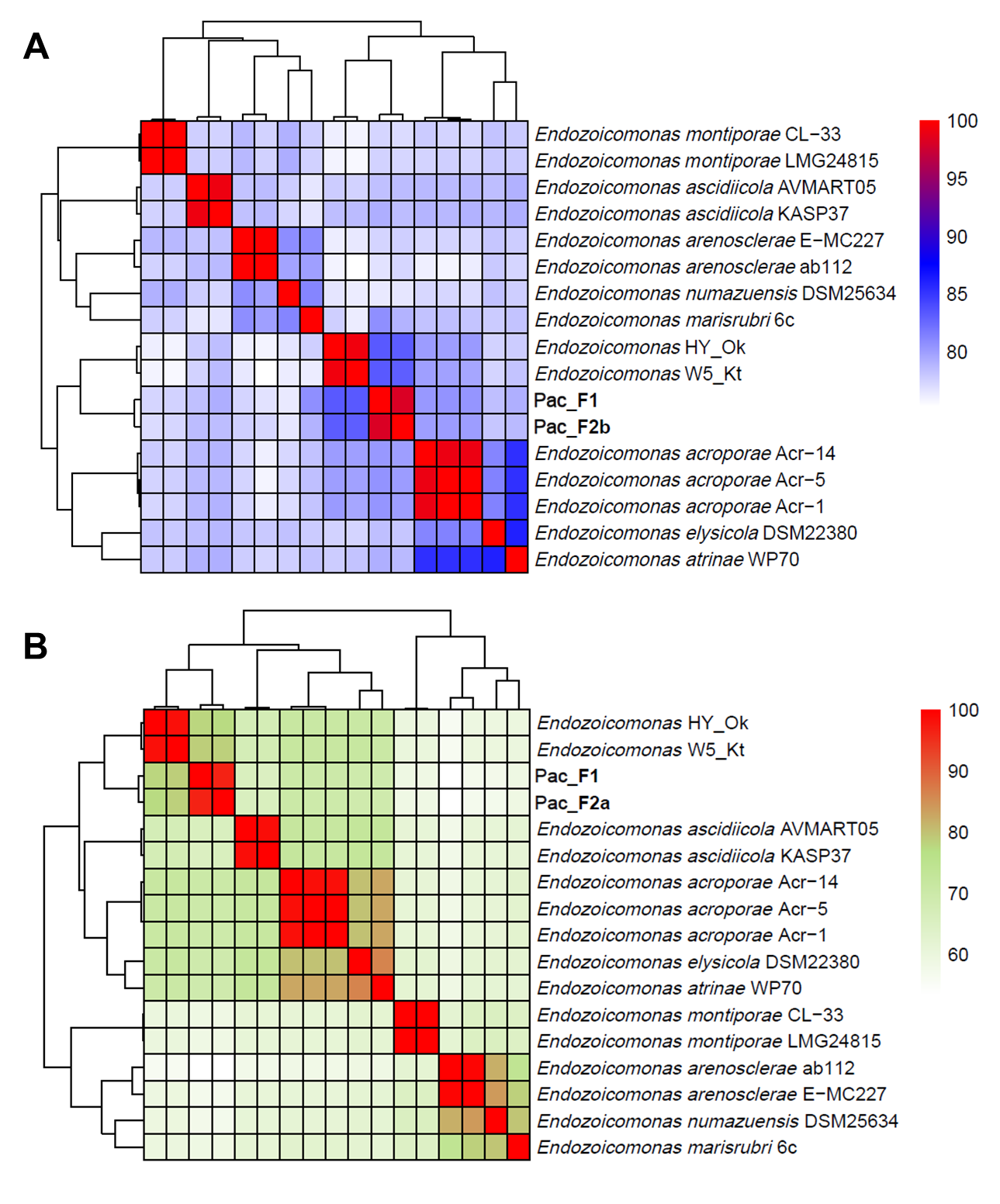

**Fig. S9:** Average nucleotide identity (ANI) (A) and average amino acid identity (AAI) (B) of Pac_F1 and Pac_F2a with other *Endozoicomonas* genomes.

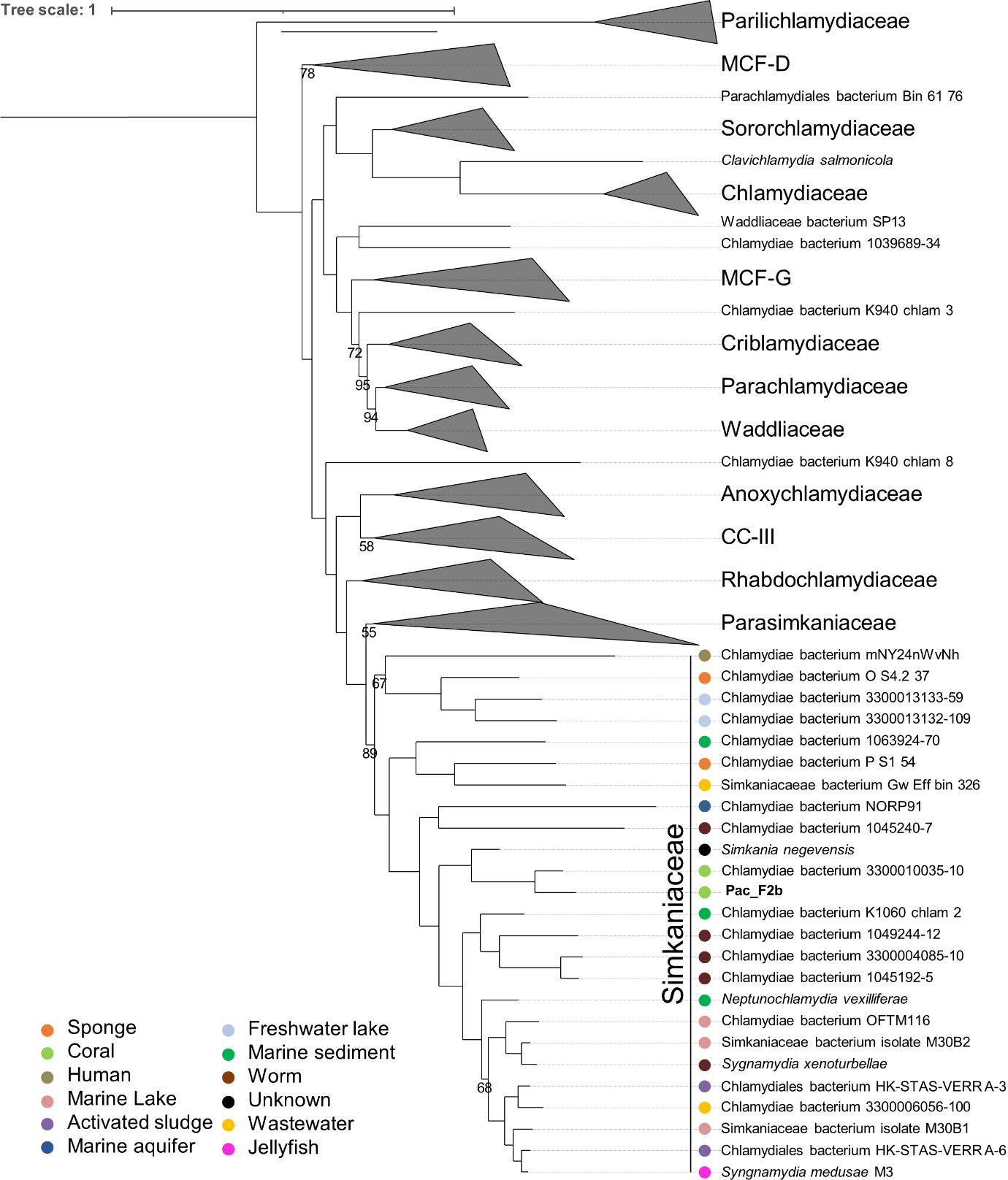

**Fig. S10:** Chlamydial maximum-likelihood phylogeny based on 15 conserved non-supervised orthologous groups (NOGs) in 139 chlamydial and 82 outgroup (Planctomycetes, Verrucomicrobia, and Lentispharae) genomes. Bootstrap support values based on 1000 replications are provided. Additional data on the reference genomes is available in Table S5B.

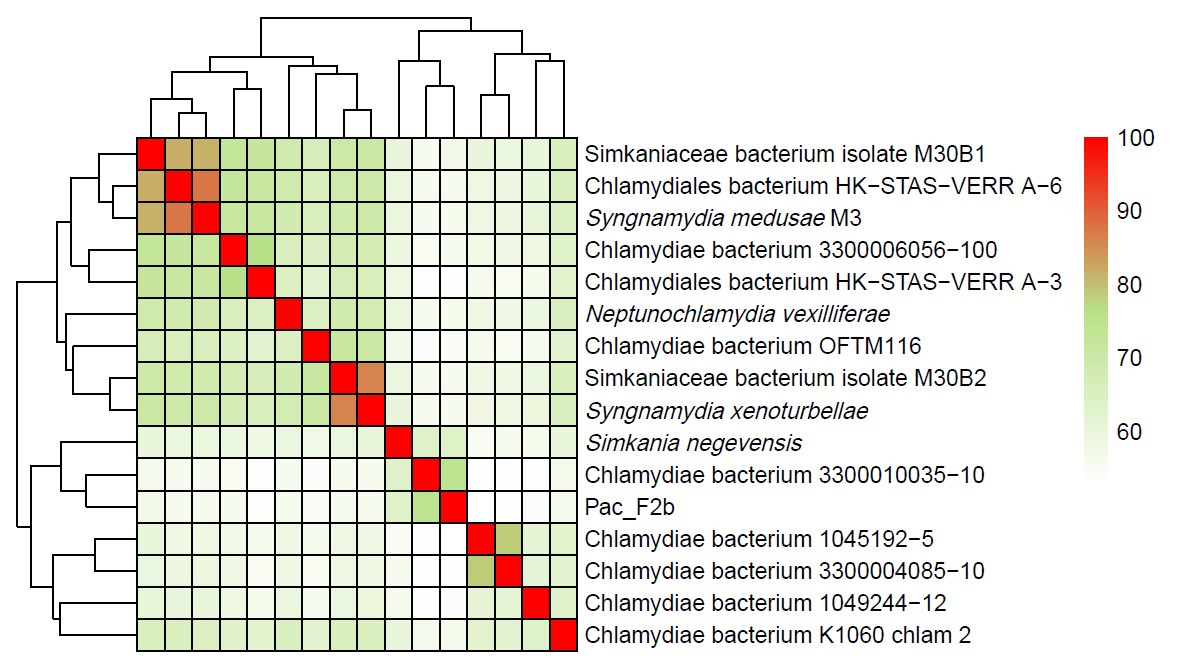

**Fig. S11:** Average amino acid identity (AAI) of Pac_F2b with other Simkaniaceae genomes.

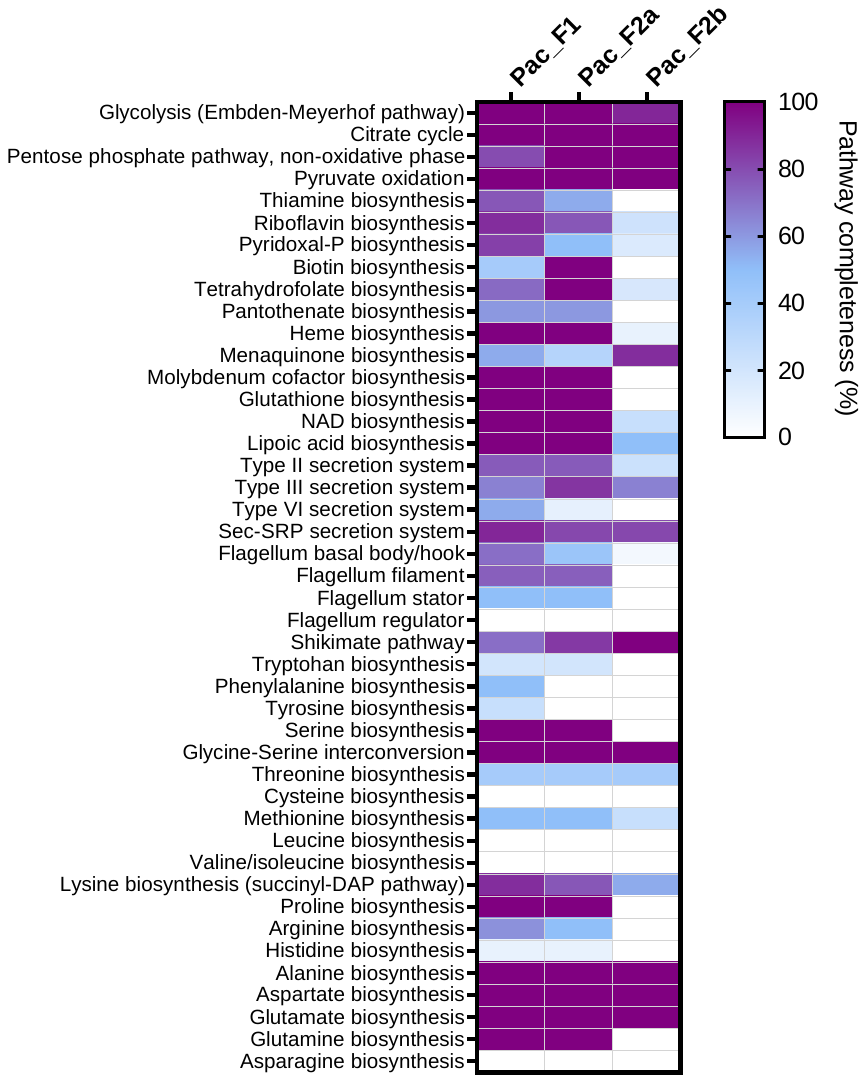

**Fig. S12:** Estimated completeness of KEGG pathways of interest in the three MAGs recovered in this study.

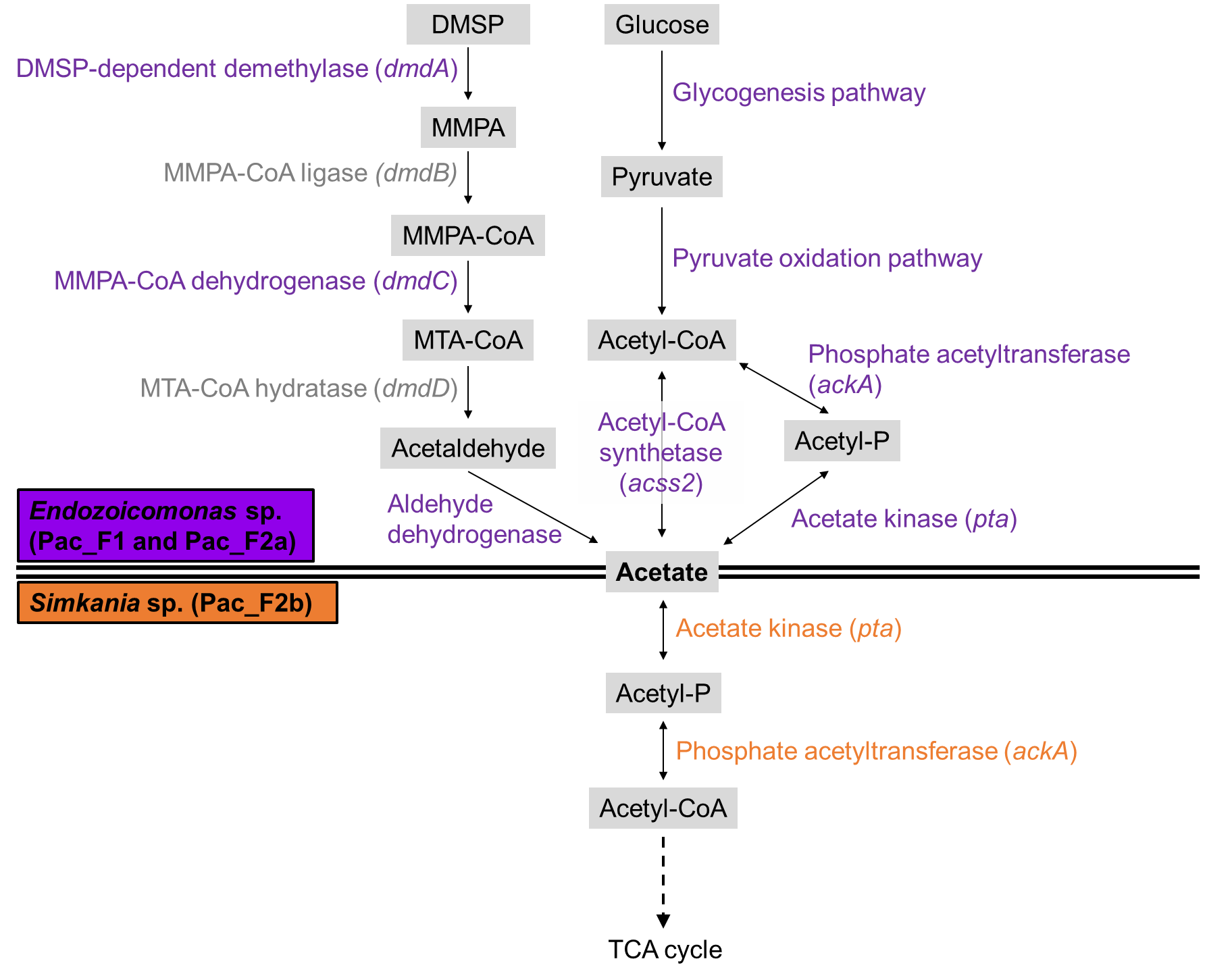

**Fig. S13**: Possible acetate cycling between *Endozoicomonas* and *Simkania*. Purple enzymes were annotated in both *Endozoicomonas* MAGs and orange enzymes were annotated in the *Simkania* MAG. Enzymes in grey font were not annotated in any MAG. DMSP: dimethylsulfonioproprionate; MMPA: methylmercaptopropionate; CoA: coenzyme A; MTA: methylthioacryloyl; TCA: tricarboxylic acid.

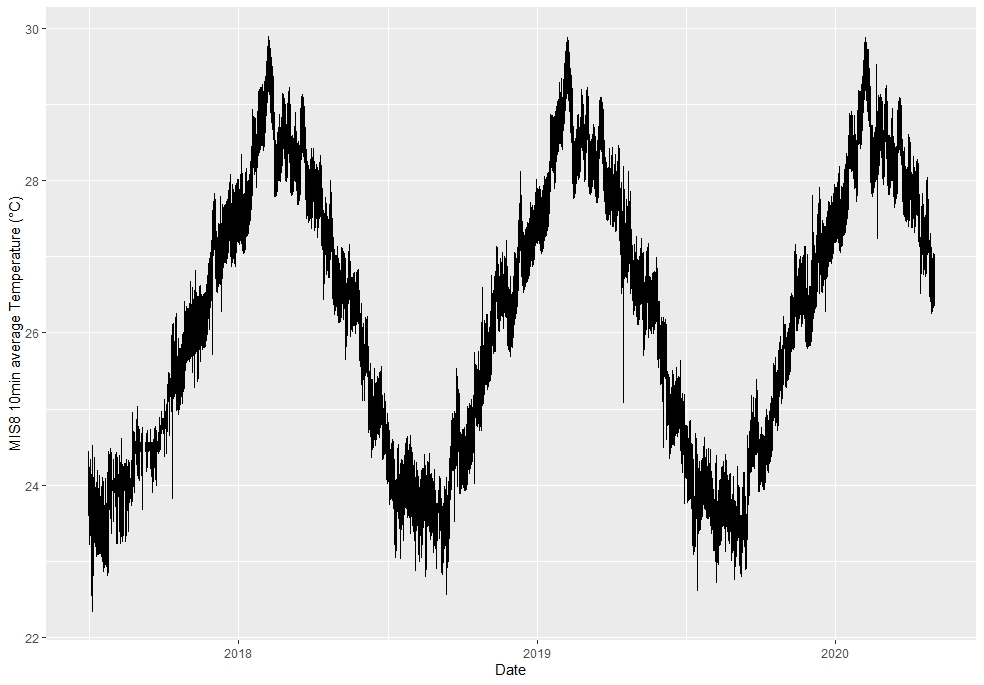

**Fig. S14:** Water temperature (10-min average) in the tank holding *P. acuta* genotypes F1_6, R2_8, and C2_12.

**Table S1:** List of *Pocillopora acuta* genotypes used in this study, including sampling location, sampling date, and which experiment they were used for. FISH: Fluorescence *in situ* Hybridization; SEM: Scanning Electron Microscopy. LCM: Laser Capture Microdissection.

(attached)

**Table S2:** Sequencing statistics for the three 16S rRNA gene metabarcoding experiments analyzed in this study. The first column (“Adult CAMAs”) refers to Figure 3A and Table S3A, the second column (“Whole larvae”) refers to Table S4, and the third column (“Larvae CAMAs”) refers to Figure 4E-F and table S3B.

| **Experiment** | **Adult CAMAs (F1­­_6 genotype)** | **Whole larvae (F1_6 genotype)** | **Larvae CAMAs (OI2 and OI3 genotypes)** |
| --- | --- | --- | --- |
| **Total Samples (negative controls)** | 17 (11) | 12 (6) | 18 (12) |
| **Raw reads** | 1642848 | 1679501 | 1323009 |
| **Reads after merging, denoising and chimera filtering** | 1147515 | 1171209 | 941134 |
| **Samples kept for analysis** | 5 | 6 | 4 |
| **ASVs after decontamination** | 16 | 179 | 18 |
| **Read per sample** | 45697 | 19075 | 13647 |
| **Contamination (%)** | 6.1 | 22.3 | 82.9 |
| **Contamination by *Brachybacterium* sp. (%)** | 5.7 | 20.4 | 77.3 |

**Table S3:** Relative abundance of bacterial ASVs in CAMAs isolated by LCM. Each column is a replicate. A: CAMAs from adults of the F1_6 genotype, generations F1 and F2. This data is summarized in Figure 3A. B: CAMAs from larvae of the OI2 and OI3 genotypes. This data is summarized in Figure 4E.

(attached)

**Table S4:** Relative abundance of bacterial ASVs in whole larvae of the F1_6 genotype. Each column is a replicate. The row highlighted in yellow is the same *Simkania* ASV observed in Figure 3A.

(attached)

**Table S5:** List of Endozoicomonadaceae (A) and chlamydiae (B) genomes used for phylogenetic analyses.

(attached)

**Table S6:** List of predicted secondary metabolites in the Pac_F1 and Pac_F2a MAGs. No predicted secondary metabolites were retrieved from the Pac_F2b MAG.

(attached)

**Table S7:** Number of eukaryotic-like protein sequences found in the three MAGs recovered in CAMA samples. Sequences were detected based on an InterProScan classification.

| Genome | **Pac_F1** | **Pac_F2a** | **Pac_F2b** |
| --- | --- | --- | --- |
| Ankyrin-repeat proteins | 114 | 104 | 0 |
| WD40 domain proteins | 4 | 3 | 0 |
| Tetratricopeptide repeat proteins | 6 | 6 | 7 |

**Table S8:** Presence of genes involved in type IV pili synthesis in *Endozoicomonas* genomes, based on a RAST analysis.

(attached)

**Table S9:** List of oligonucleotides probes used for Fluorescence *in situ* Hybridization.

(attached)

**Table S10:** Marker Non-supervised Orthologous Group (NOG) proteins used for chlamydial phylogenetic analysis.

| NOG | NOG category | NOG description |
| --- | --- | --- |
| COG0064 | J | Aspartyl-tRNA (Asn)/glutamyl-tRNA (Gln) amidotransferase subunit B |
| COG0092 | J | Ribosomal protein S3 |
| COG0233 | J | Ribosome recycling factor |
| COG0290 | J | Translation initiation factor IF-3 |
| COG0292 | J | Ribosomal protein L20 |
| COG0323 | L | DNA mismatch repair protein MutL |
| COG0335 | J | Ribosomal protein L19 |
| COG0342 | U | Preprotein translocase subunit SecD |
| COG0468 | L | DNA recombination/repair protein RecA |
| COG0532 | J | Translation initiation factor IF-2 |
| COG0536 | DL | GTPase Obg involved in cell cycle, chromosome segregation and ribosome assembly |
| COG0706 | M | Membrane protein insertase YidC |
| COG1185 | J | Polyribonucleotide nucleotidyltransferase Pnp |
| COG1530 | J | Ribonuclease G or E |
| COG1663 | M | Tetraacyldisaccharide-1-P 4’-kinase LpxK |

**Dataset S1:** Detailed Prokka and eggNOG-mapper annotations for all three MAGs sequenced in this study, Pac_F1, Pac_F2a, and Pac_F2b.

(attached)

**Dataset S2:** Orthogroup analysis of Pac_F2b against other chlamydial genomes. A: Summarized clusters of orthologous genes (COGs) present different analyzed chlamydial species set. All chlamydiae = present in at least 125 of 139 (90%) chlamydial genomes. Simkaniaceae-lineages = present in at least 32 of 35 (90%) genomes belonging to Simkaniaceae/Parasimkaniaceae/undescribed families close to Simkaniaceae. Simkaniaceae = present in at least 19 out of 21 (90%) genomes belonging to the Simkaniaceae family. Simkania = present in all three *Simkania* genomes, *i.e.* *Simkania negevensis*, Pac_F2b, and 3300010035.10. B-E: Orthologous groups (OGs), gene counts, *Simkania* refseq, and emapper annotation for all OGs present in all chlamydiae (B), *Simkania* (C), Simkaniaceae linages (D), or Simkaniaceae (E). F: Locus tags of all species in each OG (original orthofinder output). G: Number of genes from each genome belonging to each OG (original orthofinder output).

(attached)
